# Microneedle-based precision payload delivery in plants

**DOI:** 10.1101/2025.06.04.657704

**Authors:** Mingzhuo Li, Aditi Dey Poonam, Deepjyoti Singh, Jin Xu, Hongwei Jing, Ziwei Zuo, Yipeng Huang, Nana Liu, Yajun Liu, Liping Gao, Tao Xia, Anna E. Whitfield, Qingshan Wei

## Abstract

Traditional crop delivery methods, such as foliar spray and soil application, face significant limitations, including nutrient loss, environmental impacts, and low delivery efficiency. Recent advances in nanomaterials have offered novel molecular delivery platforms, but challenges such as synthesis complexity, long-term stability, and compliance with rigorous biosafety regulations persist. To provide a simpler, lower-cost, and safer alternative, we developed a polyvinyl alcohol (PVA)-based microneedle (MN) delivery system that can be precisely applied to various plant tissues (e.g., stem, lateral branch, or petiole), which demonstrates high delivery efficiency compared to the conventional methods (3.5x higher tissue accumulation) while reducing application dose (>90% less). This MN system facilitates the delivery of diverse small molecules, ranging from fluorescent dyes, growth promoters, to antiviral hormones, into plant tissues, on the other hand showing limited wounding stress to the plant. By applying fluorescent dye-loaded MNs onto tomato stems, we demonstrated effective molecular diffusion through vascular tissues. Additionally, MNs loaded with gibberellic acid (GA3) enhanced stem and branch growth in tomatoes and restored the lateral flowering phenotype in *Arabidopsis ft-10* mutants, with significant upregulation of GA receptor gene expression. Lastly, salicylic acid (SA) injections with MNs induced resistance to tomato spotted wilt virus (TSWV) in *Nicotiana benthamiana*, comparable to conventional spray and infiltration-based approaches. This easily fabricated and cost-effective MN system offers a promising tool for precision agriculture, enhancing plant health and productivity while significantly reducing the use of agrochemicals.

## Introduction

Currently, two of the most widely used methods for active delivery in agriculture are foliar spray and soil application (Niu et al., 2021; Nkebiwe et al., 2016). These two methods are popular due to their simplicity, speed, and ease of scaling up. However, with the ongoing development of the agricultural industry, significant limitations of these traditional delivery methods have become apparent (Sinha et al., 2022; Zheng et al., 2020). As larger areas of cropland and a wider variety of crops are integrated into the agricultural industry to meet the increasing food demands driven by rapid population growth, these methods of conventional compound application face new challenges (Davis et al., 2016; Fukase and Martin, 2020).

A major limitation of the foliar application method lies in its relatively low deposition efficiency and high loss of active compounds before reaching the target plant tissues. Using pesticides as an example, about 30–50% of them are dispersed into the air during the spraying process (Boonupara et al., 2023). Under unfavorable surface conditions, the deposition efficiency of foliar sprays can drop to as low as ∼20%, meaning that the majority of the applied chemicals fail to adhere to the leaf surface (Zhang et al., 2023).

While soil application saves more chemicals or nutrients, it still results in significant losses due to factors such as leaching, runoff, and volatilization. Field studies have reported that nitrogen (N) losses can range from approximately 3% to >15% of the applied amount, depending on crop type, soil properties, and management practices. For example, in a paddy field in southern China, about 3.32% of applied N was lost via surface runoff, with an additional 4.8% lost through subsurface leaching (Zeng et al., 2021). Similarly, in the Yellow River irrigation area in China, the use of conventional nitrogen fertilizers led to total N leaching loss of up to 17.7% (He et al., 2025).

Foliar application and soil-based treatment also pose significant environmental concerns when applied in high volume or under sub-optimal timing. While these methods offer convenience for crop uptake, runoff or leaching from rainfall or irrigation events can result in active compounds entering adjacent water bodies or deeper soils. For example, a new meta-analysis shows that nitrogen fertilizers (both synthetic and organic) contribute to significant nitrate leaching into groundwater, especially in coarse-textured soils and certain crop systems (Hina, 2024). Large amounts of chemicals applied to the soil can result in water contamination, soil acidification, salinization, and structural change, and eventually negative impacts to biodiversity in the soil (Alkharabsheh et al., 2021; Brichi et al., 2023; Young et al., 2021). A recent study in southwestern Germany detected large amounts of pesticides (including insecticides) in surface waters within protected areas (Schemmer et al., 2024).

In addition to generating waste and imposing environmental pressures, traditional fertilizer methods face further limitations due to the crop’s internal structure: the crop leaf cuticle, which acts as a barrier, slows down nutrient and chemical absorption (Barłóg et al., 2022; Krishnasree et al., 2021; Vasundhara and Chhabra, 2021). As such, the foliar spray approach often provides only short-term relief from nutrient deficiencies and requires frequent reapplications. Similarly, soil fertilization exhibits low efficiency, as plant uptake is delayed while nutrients move through the soil matrix into the root tissue, and it can also lead to uneven nutrient distribution (Barłóg *et al*., 2022; Dimkpa et al., 2020).

In recent years, several approaches have been developed to address these limitations in traditional crop delivery systems to make molecular delivery in plants more effective. One example of emerging technology is the use of nanomaterials as a plant delivery system (Beig et al., 2022; Jakhar et al., 2022; Liu et al., 2021; Salama et al., 2021). Nanomaterials show characteristics that protect labile compounds from fast degradation, increase the adhesion of payloads to plants, control the release of payloads, and enhance the permeability of payloads through the cuticle or internalization into plant tissues or cells (Cunningham et al., 2018; Nair et al., 2010). For example, nanocarriers (e.g., carbon nanotubes, quantum dots, and metal/metal oxide NPs) are designed to deliver DNA molecules to perform plant genome engineering, such as gene transformation, siRNA silencing, and gene editing (Cunningham *et al*., 2018; Nair *et al*., 2010; Wang et al., 2021; Zhang et al., 2021; Zhang et al., 2022; Zhang et al., 2020). However, most of these nanoscale genetic molecular delivery methods are limited by the complex nanomaterial design. In addition, the toxicity of nanoparticles to plants and animals remains a significant concern (Cunningham *et al*., 2018). Furthermore, these methods are commonly applied by spraying, which suffers from the same low uptake efficiency (∼0.1% for root application and <30% via leaf) (Tudi et al., 2021). Alternative delivery methods, such as foliar infiltration, are labor-intensive and leaf-restricted, limiting their use in the field and for broad plant species.

The microneedle (MN) is a new technique that has been applied to precision agriculture recently (Jung and Jin, 2021; Nagarkar et al., 2020). MN refers to microscopic needles, typically ranging from 150 to 1500 μm in length, 50 to 250 μm in base width, and 1 to 25 μm in tip diameter. MNs can be applied to plant tissues for the delivery of essential compounds such as nutrients, metabolites, and agrochemicals, thereby improving plant health and productivity (Ece et al., 2023; Faraji Rad, 2023). A previous example of MNs for plant delivery is biocompatible silk MNs developed for precision delivery of pesticides and other essential compounds into plants (Cao et al., 2023; Cao et al., 2020; Han et al., 2025). Even though this silk-based MN provides a new tool to deliver physiologically functional compounds into crops, the source of materials and fabrication process limit its wide application in large-scale agricultural activities. In our previous study, we developed a different MN system using water-soluble and biocompatible polyvinyl alcohol (PVA) as the base material (Paul et al., 2021; Paul et al., 2020; 2022). The PVA-based MN effectively isolated genetic molecules (e.g., DNA and RNA) from plant tissues, demonstrating its potential for use in early and in-field crop disease diagnostics (Paul *et al*., 2020; 2022). More recently, we also applied PVA MNs to crop seeds (e.g., soybeans), and demonstrated their capability for rapid seed DNA extraction for genotyping and whole genome sequencing, a much-simplified trait analysis technology that can greatly speed up the breeding process (Li et al., 2025)

In this study, we explored the capability of cost-effective and easy-to-scale-up PVA- MN to deliver small molecules into plant systems. We first tested the wounding stress induced by the PVA-MN treatment on plants and characterized the diffusion behavior of fluorescent dyes from the MN tips into tomato stem vascular tissues. Then, we demonstrated the MN-assisted delivery of plant hormone gibberellic acid (GA3) to promote plant growth and restore lateral flowering in Arabidopsis *ft-10* mutants. Finally, we tested the delivery of salicylic acid (SA) to the model plant *Nicotiana Benthamiana* to inhibit the transportation of tomato spotted wilt virus (TSWV). Together, these results demonstrated PVA- MN’s effectiveness in delivering small molecules, particularly plant hormones, to plants. This PVA- MN system holds great promise for future applications in precision agriculture.

## Results

Tomato was chosen as the model system due to its well-defined vascular structure and long petioles. Previous studies have shown that the segment between the vasculature and the epidermis ranges from 840–1040 µm for xylem and 707–925 µm for phloem in tomato plants, with xylem and phloem diameters in the tens and hundreds of micrometers, respectively (Cao *et al*., 2020). Based on these parameters, we designed and fabricated a 1000-µm-long PVA-MN as a microinjector to study payload delivery in planta (Figure 1A).

**Figure 1:**
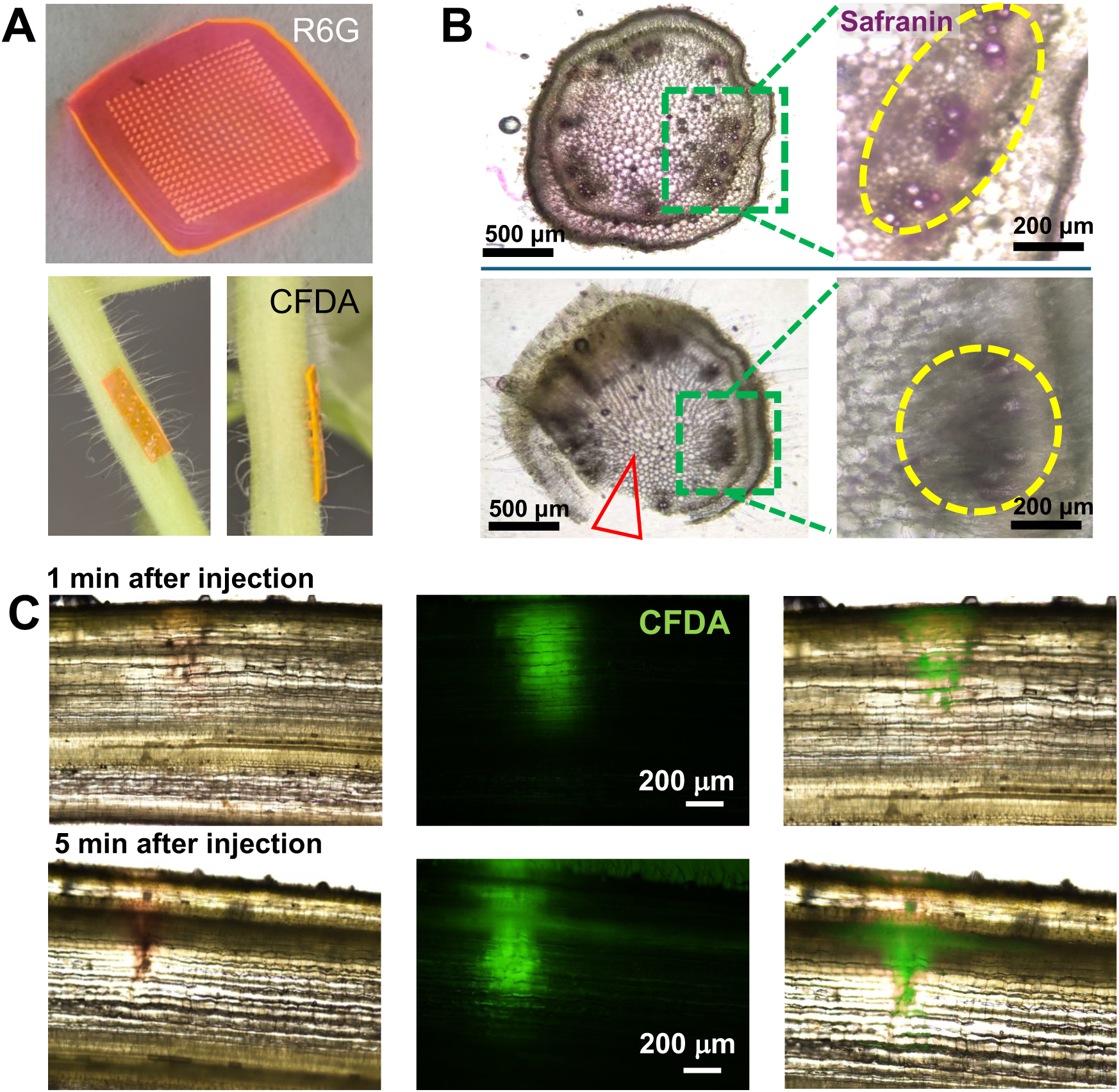
PVA-MN delivery of fluorescent dyes into tomato plant vasculature system. A) Photographs of rhodamine 6G-loaded PVA MN (top) and a PVA patch loaded with 5(6)- carboxyfluorescein diacetate (CFDA), inserted into the tomato stem (bottom). B) Cross-section microscopy images of tomato stems. Top: tomato stem without MN treatment but stained with safranin directly. The reddish cells in the zoomed-in image indicate the vascular sieve cells. Bottom: tomato stem with MN-safranin treatment (12 h). The red arrow indicates the MN penetration position; the zoomed-in image shows the accumulation of safranin dyes in the sieve cells, similar to direct staining. C) Vertical-section microscopy images (from left to right: brightfield, fluorescence, and overlap) showing CFDA delivered and transported along tomato xylem, after 1 min and 5 min of PVA MN injection.

### Effect of MN-induced mechanical stress on plant growth

PVA is a synthetic polymer with several distinct characteristics, such as water solubility, biodegradability, non-toxicity, and tunable mechanical properties, making it a promising candidate for applications in plant delivery. To test whether PVA MN could be used for chemical delivery in plant tissue, we first examined its mechanical damage to plant health. Due to the strong hygroscopic nature of PVA, we observed that leaf tissues punctured by the PVA MN dehydrated rapidly if the MN was not removed immediately after penetration (e.g., contact time >20 s), causing surrounding tissues to wilt within an hour post-puncture (Figure S1). In contrast, when the microneedle was applied to the stem, although localized dehydration occurred at the puncture site, the vascular bundles remained largely unaffected and continued to function properly (Figure S1). These results indicate that the microneedle contact time for leaf delivery needs to be controlled, whereas the stem is a more robust delivery site for PVA MN.

The morphology changes of MN tips before and after stem penetration were characterized by scanning electron microscopy (SEM) (Figure S2A,B). We found that after 30 seconds inside the stem, the MN tips were softened, and some needle tips were partially dissolved by the cellular fluids within the stem. The PVA-MN weight loss was also monitored. Before MN punching on the tomato stem, the average weight of a small PVA-MN patch (9 x 6 array) is around 2.5 mg (Figure S2C). After MN injection, the average weight of used MN is about 7, 10, or 13% lower than that of the new MN patches with contact time of 1, 2, and 15 min, respectively, indicating PVA material dissolving over time during the MN treatment process (Figure S2C). Additionally, we tested the mechanical properties of PVA MN under compression on the stem. The results showed that the average needle tip break force for tomato stem penetration is about 0.2N (Figure S2D). These observations indicate that PVA MN exhibits an excellent combination of rapid water solubility and mechanical strength for plant delivery.

Next, we investigated whether the PVA MN punch at the stem would have a negative impact on plant health and growth over time. We examined the damage caused by MN insertion on tomato stems under a microscope (Figure S3). On the punctured surface, the tomato stems appeared slightly hardened, but no visible oxidation or darkening was observed at the puncture sites. This phenomenon could be attributed to PVA’s water-absorbing property, which causes local cell dehydration and the formation of a protective layer. The microscopic images of the tomato stem cross-section showed that the pith region at the center of the stem developed a few void zones at 2 mins after MN punch, indicating acute cell damage (Figure S3B, middle column). However, over time, the pith cells repaired themselves and filled these gaps within 48 hours after MN puncturing (Figure S3B, right column). Meanwhile, the vascular bundle cells at the puncture site formed a protective layer upon cell death, which helped prevent further internal damage. In addition, no excessive tissue oxidation or darkening was observed around the puncture site externally (Figure S3, top row).

The tomato *SIBOH1*(Respiratory burst oxidase homolog) gene is a commonly used indicator of the abiotic stresses. We analysed the *SIBOH1* gene expression in the stem after the PVA MN punch. The results showed that the transcription level of *SIBOH1* was induced by 2-6 fold immediately after MN treatment (30 mins) (Figure S4A). However, the *SIBOH1* expression decreased to a normal level after 48 h of MN punch (Figure S4A). At the same time, we also tracked the *TomLOXD* gene, which is a classical lipoxygenase gene induced by mechanical damage. Gene expression analysis showed that the relative expression level of *TomLOXD* was also induced 3-4 fold after 30 min of MN wounding; after 48 h, the relative expression of *TomLOXD* was down regulated to the normal range (Figure S4B). These molecular results match the previous microscopic observations (Figure S3), suggesting that the MN-treated plant stem showed a certain level of acute mechanical damage, which was eventually self-recovered.

Meanwhile, we compared the growth of wild-type (WT) tomato plants (without MN punch) with those that had been subjected to three rounds of PVA-MN puncturing. Phenotypically, there are no significant differences in growth height between WT and MN-treated plants (Figure S5A). Quantitative measurements of plant height were taken at 1, 6, and 15 days after puncturing and revealed no noticeable differences compared with WT plants (Figure S5B). Together, these findings (Figure S3-5) suggest that PVA MN puncturing on tomato plant stems, even with repeated treatments, does not exert a significant inhibitory effect on normal plant growth.

### Delivery of fluorescent molecules in plant tissue

Fluorescent or colorimetric dyes were chosen as the first model small molecules for MN-assisted plant delivery. Because of the water-dissolving characteristic of the PVA material, we speculated that the small molecules are more likely to be transported by xylem. Safranin is a common plant histological staining dye for xylem ducts. We loaded safranin into the PVA- MN at a concentration of 1%. After stem injection, we observed a faint reddish color accumulation around the vessel cells after MN injection, suggesting that the MN-delivered dyes were mainly transported through the xylem system after MN punching (Figure 1B).

We also incorporated fluorescent dyes, such as rhodamine 6G (maximum absorption at 526 nm) and 5(6)-carboxyfluorescein diacetate (CFDA, maximum absorption at 492 nm), into the PVA-MN. We observed the diffusion of CFDA into surrounding tissues within 5 minutes of PVA- MN injection (Figure 1C). To analyze molecular transport in the xylem, we performed tangential sectioning of the stem and monitored the dye diffusion at 1-, 5-, and 10-min post-injection by fluorescence imaging. The results showed a significantly more widespread distribution at 5 min post-injection than at 1 min (Figure 1C). Quantification of relative fluorescence intensity showed a 32% and 77% increase of dye diffusion area in stems at 5- and 10-min post-injection, respectively (Figure S6). Additionally, the chemical dyes were observed to travel along the xylem flow in the tomato stem (Figure S7). These findings strongly suggest that the PVA-MN tip effectively reaches the vasculature and that chemical compounds embedded in the PVA-MN can diffuse and be transported along the vascular, especially the vessel cell delivery system in tomato plants.

### Delivery of growth regulator GA3 to promote tomato plant growth

To demonstrate the utility of PVA MN to deliver plant growth regulators (PGRs) in crops, we investigated the delivery of gibberellic acid (GA3) to tomato plants. GA3 is a plant hormone that plays a crucial role in regulating various growth and developmental processes in plants, such as seed germination, stem elongation, flowering, and fruit development (Gupta and Chakrabarty, 2013). In the current study, the tomato stem was selected as the injection site.

Twenty-five-day-old tomato plants (n=12) were injected with the PVA-MN on the stem tissue (Figure 2A, Figure S8). Following the initial MN injection, secondary and tertiary injections were performed 3 and 6 days later, respectively. Plants treated with PVA MN without GA3 (negative control, or “Control-MN”), plants with direct GA3 injection at the stem by a syringe (“GA3-Injection”), and plants sprayed with GA3 on their leaves (positive control, or “GA3-Spary”) were included in the experiment (Figure 2A). In addition, healthy plants without any microneedle or GA3 treatment were used as the wild type (“WT”, Figure 2A). The PVA-MN patch contains 500 μM GA3, which is higher than the loading concentration of GA3 in the spray method (100 μM). At the same time, for direct stem injection, we used 100 μL of GA3 solution with a concentration of 100 μM. However, due to the much smaller delivery volume of the PVA- MN method (only an array of ∼ 4 x 10 needles penetrated stems) compared to 5-10 mL of solutions applied in the spray method, three treatments of PVA-MN deliver an estimated total amount of 0.05-0.075 μmol of GA3, which is only about 10% of the spray method, and 50% of the stem-based direct injection method (Table1).

**Figure 2.**
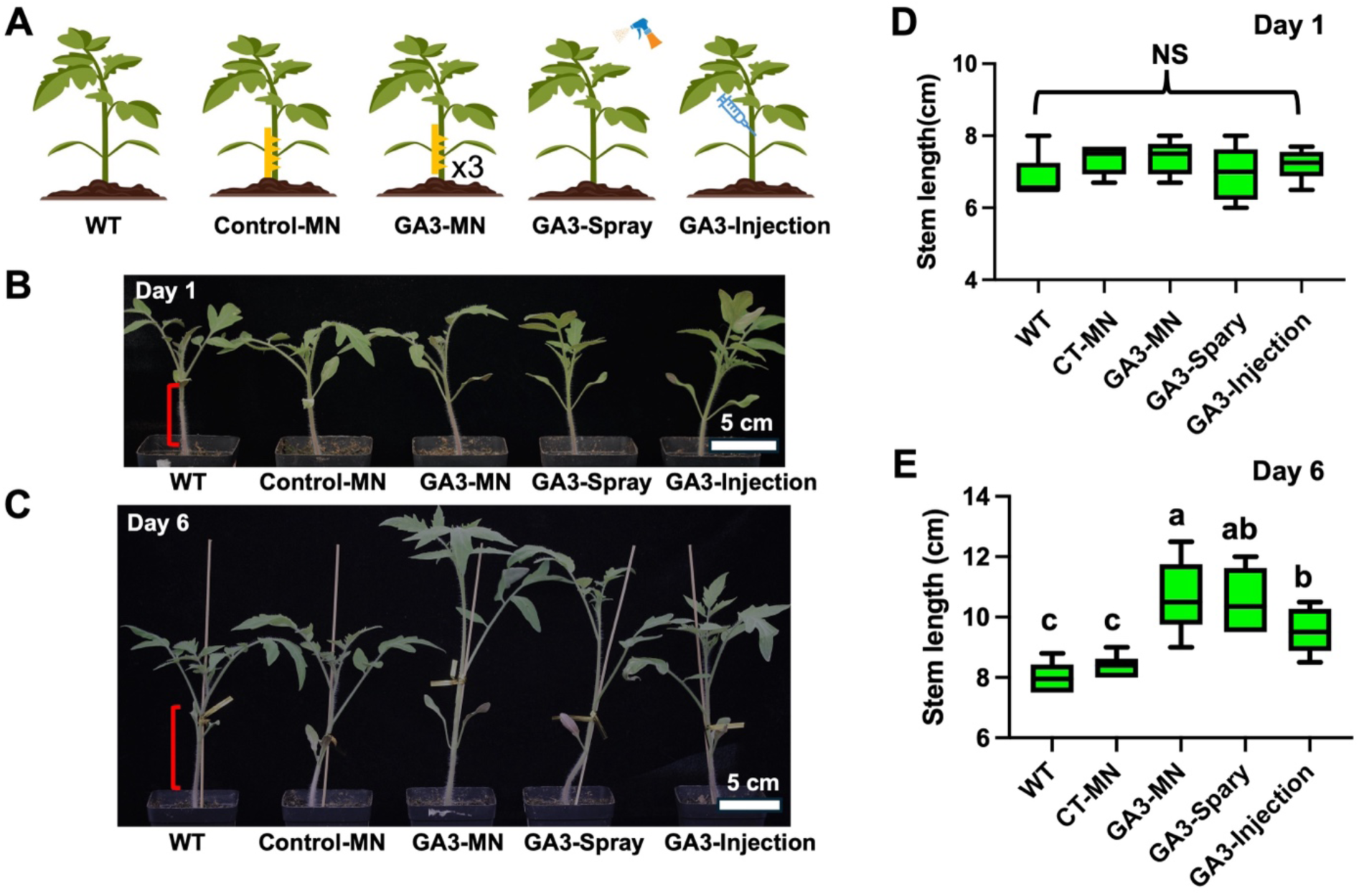
**Tomato growth phenotype under different GA3 treatments**. A) Schematic of different GA3 treatment methods on 25-day-old tomato plants. B,C) The growth phenotype of tomato plants at Day 1 and Day 6, respectively. D) The stem length of the tomato plants at Day 1 under different GA3 stimulations (n=12). NS = not significant. E) The stem length of the tomato plants at Day 6 under different GA3 stimulations (n=12). Different letters above the bars indicate statistically significant differences between groups (P < 0.05). Groups that share a letter are not significantly different (P > 0.05). CT = control.

After six days, we observed that tomato plants treated with GA3, including GA3-MN, GA3-Spary, and GA3-Injection, exhibited a faster growth rate compared to control plants (both WT and Control-MN, Figure 2B, C, E). The stem lengths (labeled in red, Figure 2B, C) were quantitatively measured at the same time, revealing that after 3 days, the heights of GA3-MN and GA3-Spray-treated tomato plants were 18%–21% higher than those of the WT or Control-MN plants (Figure S8). At the same time, the GA3-Injection plants showed an increase of around 10% in height (Figure S8), which was less effective than the GA3-MN or GA3-Spray groups. After 6 days, GA3-treated (including the GA3-MN, GA3-Spray, and GA3-Injection) plants showed an increase of 20%–43% in height compared to the non-treated controls (Figure 2E). The GA3- Injection group is again slightly less effective than GA3-MN or GA3-Spray. A similar trend was also observed on Day 9 (Figure S8B) and Day 15 (Figure S9). In particular, on Day 15, the GA3- MN group eventually outgrew all other control groups (Figure S9). In our test, direct injection of GA3 into the tomato stem by a syringe did not promote plant growth to the same extent as leaf spray or MN delivery. This is probably due to the fact that the direct stem injection method is very hard to control, where a precision control of injection rate is required to avoid liquid overflow. In contrast, the MN approach is much easier to apply without the need for flow rate control and therefore is more suitable for field or greenhouse applications.

We also tested the effect of the application frequency of GA3-MN treatment on tomato growth (Figure S10). The tomato plants for the MN injection experiment were divided into three groups, which received one (1x), two (2x), and three times (3x) of injections, respectively. All plants were injected on the first day; then, the second and third groups were injected again after three days; only the third group received a final injection after six days. Plant height was measured simultaneously throughout the experiment. The results showed that for GA3 delivery, one MN treatment is sufficient to promote tomato growth (Figure S10). On Day 6 and Day 9, the plant height in group 2 and group 3 with more injections revealed larger variations compared to group 1 with only 1x MN injection (Figure S10B). These results demonstrated that for GA3 delivery by PVA- MN, the application frequency can be as low as 1x.

Furthermore, GA3 delivery by PVA MN could also be performed on the lateral branch by using 40-day-old tomato plants (Figure S11A and B). The injection process is the same as the stem treatment. Due to the difference in the initial length of each lateral branch, we calculated the relative lateral branch elongation to evaluate the GA3 stimulation. The results showed that after 3 days of treatment, the average lateral branch of GA3-MN and GA3-Spray-treated tomato plants grew around 5% more than the WT and Control-MN plants (Figure S11C); after 6 days post-injection, the average lateral branch length of the GA3-MN group increased by nearly 10% of the control groups (Figure S11D).

In addition to the stem or branch growth length difference, we observed that the tomato plants in GA3-treated groups (GA3-MN or GA3-Spray) showed a noticeable color difference from the 4th to 6th compound leaf (newly emerging leaves) (Figure S12A). We assume that this phenotype difference is probably due to the accelerated growth triggered by GA3. The concentrations of chlorophyll a (p = 0.0049) and b (p = 0.0025) in the 4th to 6th leaves (mixed samples) were also quantified. The results showed that in GA3-treated tomato, the leaf chlorophyll a and b content decreased by 6% to 10% compared with WT and Control-MN plants (Figure S12B and C), suggesting a faster growth rate in GA3-treated groups.

To investigate whether the GA3-MN treatment would trigger the intrinsic gene transcription level changes in tomato plants. We conducted a series of reverse transcription-quantitative polymerase chain reaction (RT-qPCR) experiments using tomato leaf samples after different treatments. We selected the GA signal receptor, two tomato *GID* genes (Shohat et al., 2021), as targets and ran gene expression analysis. The results revealed that after GA3-MN or GA3-Spray treatments, both *SiGID1* and *SiGID1b1* genes showed significant increases in transcription level by 2 to 15-fold compared to the WT and Control-MN plants (Figure 3). The expression level of GA3-MN group is generally comparable to the GA3-Spray group (Figure 3). Interestingly, in individual plant #9, the MN treatment promoted higher *GID* gene expression level than conventional spraying. All the above data strongly suggest that the PVA-MN-GA3 injection promotes the stem or branch growth similarly to the foliar spray method, while reducing the application dose by 10∼20 times. Consistently, the gene expression involved in the GA3 signal in tomato plants was also upregulated, confirming the effectiveness of the delivery of GA3 by the MN patch into tomato plants.

**Figure 3.**
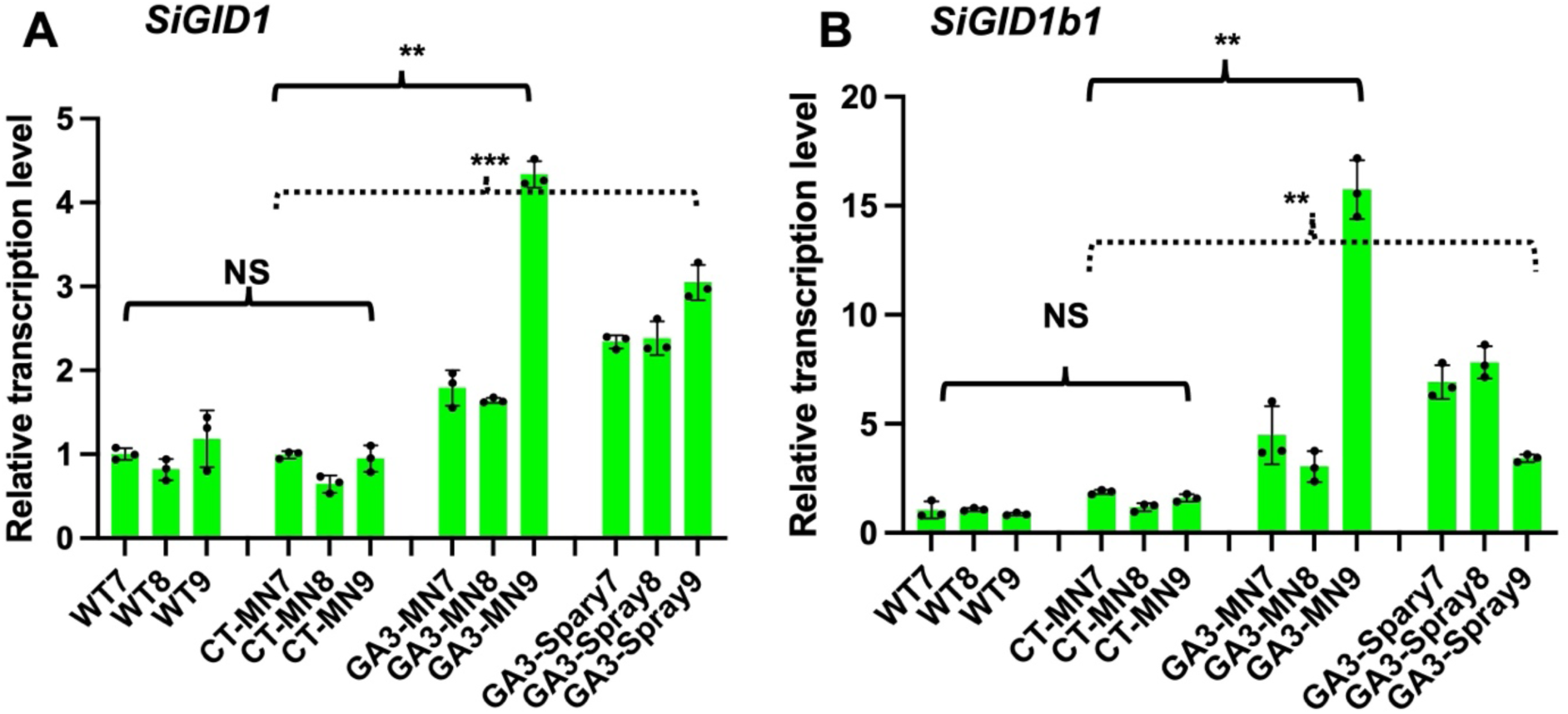
Tomato gene expression levels under different GA3 stimulations. A) The relative transcription level change of *SiGID1* genes under the different GA3 treatments in tomato plants. Each group has three different plant lines (plants 7, 8, 9). B) The relative transcription level change of *SiGID1b1* genes under the different GA3 treatments in tomato plants. Each group has three different plant lines (plants 7, 8, 9). CT = control. NS = not significant, *P < 0.05, **P < 0.01, ***P < 0.001.

### PVA-MN for GA3 Delivery to *Arabidopsis ft-10 mutant*

Besides tomatoes, we also used a model plant *Arabidopsis thaliana* (*A. thaliana*) and investigated the PVA- MN injection effects on the GA3 signaling process. *A. thaliana* showed multiple advantages in phenotyping analysis such as well-characterized biological activities (including development, metabolism, hormone signaling, and stress responses), ease of genetic manipulation, and comprehensive genomic information.

Numerous studies have shown that applying gibberellins (such as GA3) in *A. thaliana* promotes the termination of vegetative growth, regulates flower formation, accelerates bolting, and results in plants with fewer rosette leaves and more cauline leaves (Bao et al., 2020). Under normal conditions, wild-type *A. thaliana* Col-0 plants produce around 14 rosette leaves during the vegetative growth phase before inflorescence emergence at approximately 23 days under a 16- hour light regime. Some studies report 11–13 rosette leaves between days 21–24 (Cao *et al*., 2023). Cao and colleagues studied the *Arabidopsis ft-10* mutant, a *FT* gene loss-of-function mutant with a clear late-flowering phenotype under long-day conditions, to evaluate their silk-based microneedle delivery system (Cao *et al*., 2023). This mutant disrupts non-GA3-related pathways that predominantly regulate flowering under long-day conditions (Figure S13A). Unlike the previous silk-based microneedles, our study applied PVA-MN to the petiole of a single rosette leaf in *Arabidopsis* plants (Figure S13B, C). The PVA- MN used in this experiment was 800 µm in length.

To assess the effectiveness of PVA-MN in delivering GA3 to *Arabidopsis* plants, we again compared MN-mediated delivery with foliar spray. Additionally, an empty PVA-MN (without GA3) was used as a negative control (Control-MN, Figure 4A). GA3 was delivered via PVA-MN injections at multiple leaves every 3 days, for a total of three applications (x3), similar to the frequency of foliar sprays. For MN injection on *Arabidopsis* vein, a small array of 2 x 10 needles was used. The leaf spray method used 2∼4 mL of GA3 solution instead. The total GA3 amount delivered by the spray method is nearly 10 times that used in the MN injection methods (Table1). We observed that both GA3-Spray and GA3-MN treatments resulted in an early bolting phenotype compared to the *ft-10* mutant control and the Control-MN group (MN without GA3) (Figure 4B). Although GA3-MN-treated plants bolted earlier than the controls, their response was slower than that of GA3-Spray-treated plants. It is because *Arabidopsis* exhibits a distinct, two-stage developmental pattern, in which vegetative growth precedes reproductive growth. Initially, the plant dedicates its energy to forming a rosette. The main stem’s internodes remain compressed during this entire phase. Once it receives the necessary environmental and internal cues, the plant undergoes a critical phase transition that initiates rapid, dramatic elongation of the central stem, known as bolting. In our experiment, the PVA-MN injection was performed at the petiole site, which is different from the previous injection into the stem of tomato plants. The diffusion of GA3 to the whole plant from the petiole site is likely slower than that injected directly into the stem.

**Figure 4:**
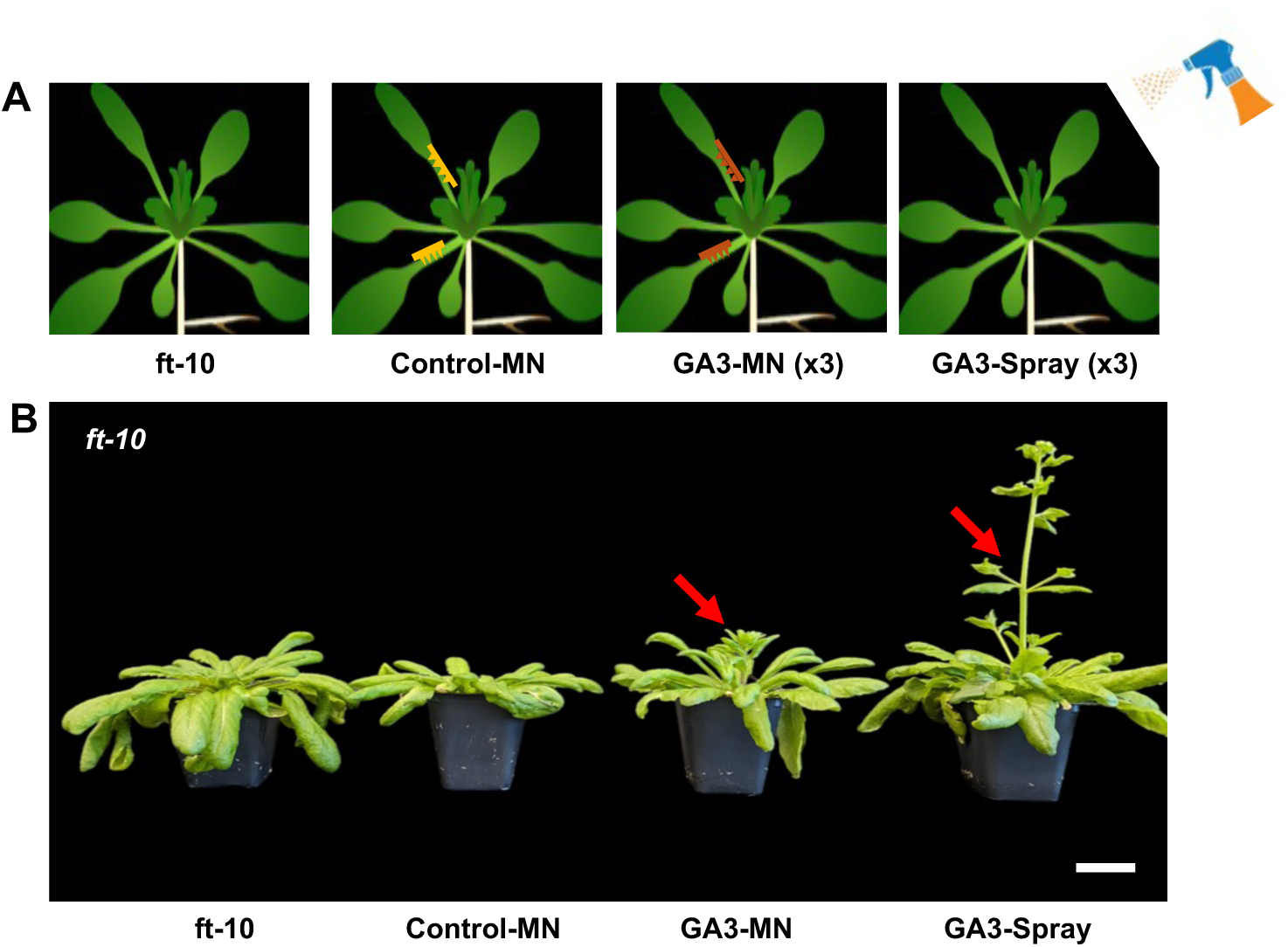
PVA-MN for GA3 delivery to *Arabidopsis ft-10* mutant. A) Schematic of different GA3 delivery methods to the *Arabidopsis ft-10* mutant. B) Representative image of *ft-10* plants 10 days after different GA3 treatments. MN injection and spray were carried out every three days. 4 groups of *ft-10* (n=5 for each group) were treated differently (including *ft-10*, Control-MN, GA3- MN, and GA3-Spray).

Furthermore, for MN treatment, we injected only three out of more than twenty inflorescences, meaning the total GA3 applied in the GA3-MN group was significantly lower than that in the GA3-Spray group. All these may contribute to the different flowering time observed between the GA3-MN and GA3-Spray groups.

In addition, rosette and cauline leaf numbers were also recorded throughout the growth period. GA3-MN injections produced phenotypes in *ft-10* similar to those observed in GA3-Spray plants, with changes in rosette and cauline leaf numbers and leaf size (Figure 5A). These phenotypes were significantly different from those *ft-10* mutants untreated with GA3 (Figure 5A). This confirms that GA3 delivered via microneedles effectively altered plant metabolism. Interestingly, Control-MN plants (without GA3) produced fewer rosette leaves than the untreated *ft-10* mutant, suggesting that PVA- MN injection may induce some stress even in the absence of GA3 (Figure 5A). These phenotypic findings demonstrate that PVA-MN is effective in delivering GA3 to *Arabidopsis* plants and changes their growth behavior accordingly.

**Figure 5.**
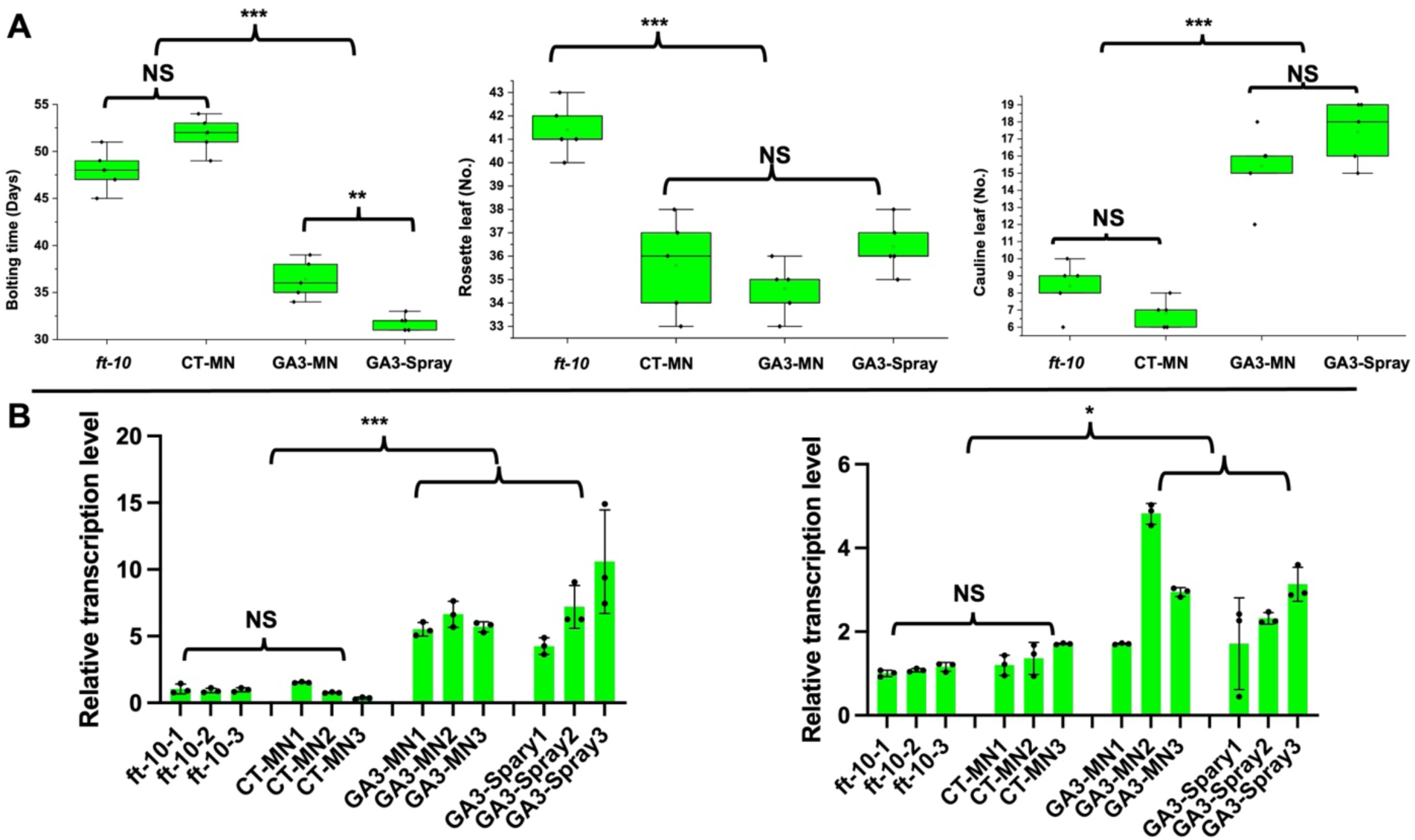
Different GA3 delivery treatments alter *Arabidopsis* phenotype and gene expression in the GA3 signaling pathway. A) The bolting time, rosette leaf number, and cauline leaf number change under different GA3 treatments in *Arabidopsis* ft-10 mutants. B) The relative transcription level change of *AtGID1b* and *AtGID1c* genes under the different GA3 treatments in *Arabidopsis* plant. Each group has three different plant lines (plants 1, 2, 3). CT = control; NS = not significant; *P < 0.05, **P < 0.01, ***P < 0.001.

In *Arabidopsis* plants, there are three GA receptor isoform genes, namely *AtGID1a*, *1b,* and *1c*. Using RT-qPCR, we analyzed transcript levels of these three genes with and without the GA3-MN treatment (Figure 5B). We detected that the *AtGID1a* showed a low transcript level in the *Arabidopsis ft-10* mutant background, but the other two GA receptor genes, the *AtGID1b* and *AtGID1c*, showed a significant increase in the mRNA level by 2 to 10 fold after the GA3-MN and GA3-Spray treatments (Figure 5B). These molecular results confirm the previous phenotypic observations, suggesting that PVA-MN is an effective tool for delivering plant hormones in *Arabidopsis*.

### Delivery of SA for induced TSWV resistance in *Nicotiana benthamiana* plant

To test whether plant hormones can be successfully delivered into plants by using the MN, we selected salicylic acid (SA) as a testing model and fabricated a PVA-MN patch loaded with 5 mM SA. SA is an important plant hormone that plays a significant role in triggering the pathogen resistance response. To measure SA delivery efficiency, we used a 3 x 18 MN array applied to the *Nicotiana Benthamiana* petiole with 2 min of contact time (“SA-MN”). At the same time, a 500 mM SA solution was sprayed on the leaf surface (“SA-Spray”) or infiltrated into the leaf (“SA- Infiltration”) as the control (Figure 6A). Each *Nicotiana Benthamiana* plant was sprayed with 5- 10 mL of SA solution, and the injection volume for the SA-Infiltration method is approximately 1 mL. After 24 h of treatment, 100 mg of leaf tissue was sampled and ground in liquid nitrogen. Plant tissue SA was isolated by an extraction buffer and analyzed by high-performance liquid chromatography combined with a mass spectrometer (HPLC-MS).

**Figure 6.**
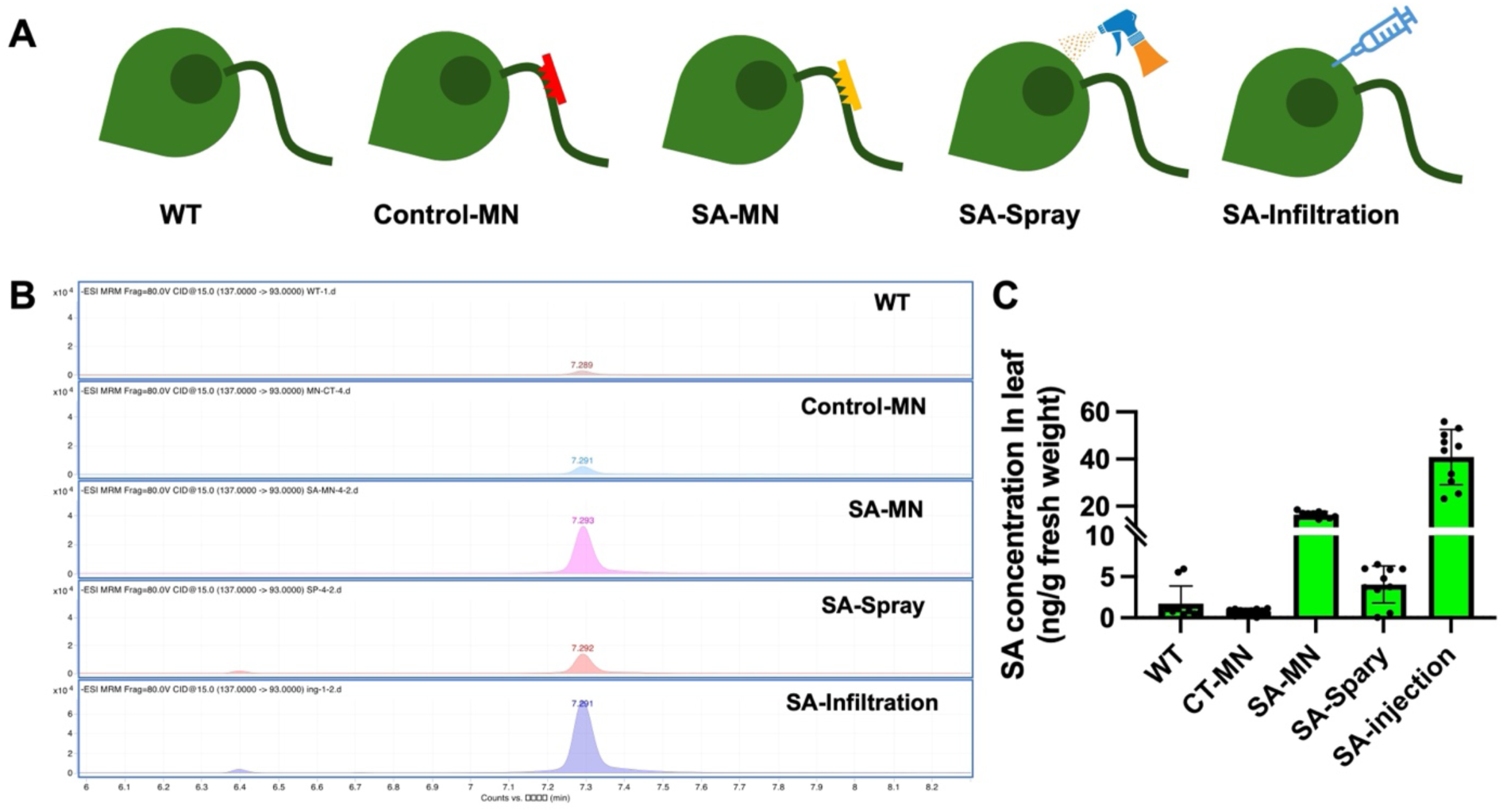
PVA-MN delivery of SA in *Nicotiana Benthamiana*. A) Schematic illustration of different SA delivery methods to *Nicotiana Benthamiana* plant on the petiole; Later, the dark region on the leaf was sampled for SA concentration analysis. B) HPLC-MS analysis showed the presence of SA (7.29 min) in leaf tissues with different delivery methods. C) The SA concentration in different samples showed that PVA-MN has a higher delivery efficiency than the spray method. (n=10). CT = control.

The HPLC-MS/MS results showed that in WT and Control-MN plant leaves, there was very little SA (Figure 6B, Figure S14). In contrast, the leaves in the SA-MN, SA-Spray, and SA- Infiltration groups all showed relatively high SA concentrations (Figure 6B, Figure S14A). Notably, the SA-MN group showed a higher peak than the SA-Spray method (Figure 6B), indicating a higher delivery efficiency for the SA-MN method. We also quantified the SA concentrations in different samples. The results showed that leaves with SA-Infiltration had the highest SA content at 40 ng/g (Figure 6C). On the other hand, the MN-treated leaves accumulated a SA concentration of 14 ng/g, while the spray method only showed a SA concentration of 4 ng/g, the lowest among the three (Figure 6C). Overall, these results verify that the PVA-MN approach demonstrated a high delivery efficiency despite its low loading capacity.

Tomato spotted wilt virus (TSWV) is a plant virus that belongs to the family *Tospoviridae* and is one of the most economically significant plant viruses affecting crops worldwide (Best, 1968). The infected plants often show necrotic spots, wilting, chlorosis (yellowing), stunted growth, leaf deformation phenotype, and in some cases, ring spots or concentric patterns on leaves and fruits. These symptoms lead to reduced yield and poor fruit quality. Plant hormone SA has been shown to play a significant role in triggering the resistance response to TSWV in plants (Turina et al., 2016). SA is a crucial plant hormone involved in systemic acquired resistance (SAR), a defense mechanism that provides broad-spectrum resistance to a variety of pathogens, including viruses like TSWV (Durner et al., 1997). In the following experiment, we used PVA-MN to deliver SA directly into plant tissues and investigated whether the MN microinjection could help the model plant *Nicotiana Benthamiana* generate resistance to TSWV. Normally, the virus would move within plants through a well-coordinated process that involves both cell-to-cell movement and systemic movement via the plant’s vascular system. To monitor TSWV movement in the plant, TSWV infectious clone generated by Liu et al was engineered, and Cas9 in the infectious clone was replaced with RUBY reporter (Liu et al., 2023), which makes the viral movement visible to the naked eye (Figure S15).

We compared the SA delivery and treatment efficiency of four different methods, including Control (no SA delivery), SA- MN, SA-Spray, and SA-Infiltration (Figure 7A). For SA- MN, the MN patch loaded with SA was applied to *Nicotiana Benthamiana* petiole for 4 times in total (1x before and 3x after TSWV inoculation). The SA-Spray method was performed 2 times (before and after inoculation), and the SA-Infiltration method was only performed once on *N. Benthamiana* (Figure 7B). For PVA-MN injection, a small MN array of 4x10 needles was used. The spray and infiltration treatments used 20-25 mL and 8-10 mL of SA solutions, respectively. The absolute amounts of SA delivered to tobacco plants by different delivery methods are summarized in Table 1.

**Figure 7.**
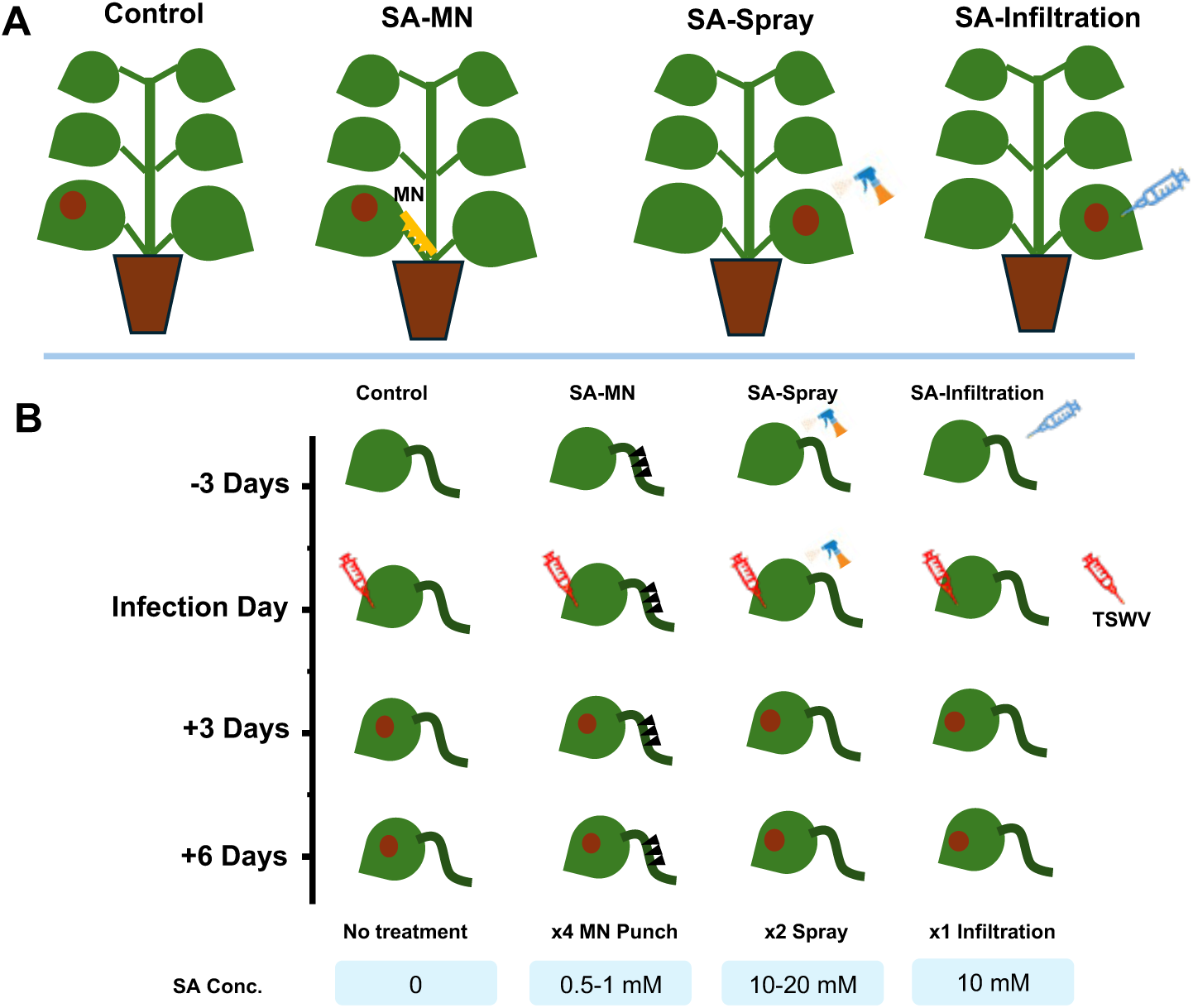
SA delivery to prevent TSWV virus movement in *Nicotiana benthamiana* plants. A) A schematic showing different SA delivery approaches. B) A treatment timeline showing the SA treatment process for SA-MN, SA-Spray, and SA-Infiltration methods in *Nicotiana benthamiana* plants, as well as the concentrations of SA used in different delivery approaches.

**Table 1.** Comparison of plant hormone consumption in different delivery methods

| Plants | Cargo | Delivery Methods |  |  |
| --- | --- | --- | --- | --- |
| Tomato | GA3 | GA3-MN | GA3-Spary | GA3-Stem Injection |
| | | 0.05~0.075 $\mu\text{mol}$ | 0.5~1 $\mu\text{mol}$ | 1~1.5 $\mu\text{mol}$ |
| *Arabidopsis* | GA3 | 0.025~0.03 $\mu\text{mol}$ | 0.2~0.4 $\mu\text{mol}$ | |
| *Nicotiana benthamiana* | SA | SA-MN | SA-Spray | SA-Leaf Infiltration |
| | | 1.2~1.6 $\mu\text{mol}$ | 200~250 $\mu\text{mol}$ | 50 $\mu\text{mol}$ |

In the first trial, the SA-MN group contained 2 mM SA, while the SA-Spray and SA- Infiltration groups used only 500 μM SA, considering their larger application volumes. Still, the estimated amount of SA delivered to the plants by the MN approach is only about 0.5-1% of the spray or infiltration method (Table 1). Four *Nicotiana Benthamiana* plants were used for each treatment, and three leaves were randomly selected from each plant for viral inoculation. During the infection process, the gradual deepening of RUBY pigmentation was observed, indicating the progression of viral infection. On the third- and sixth-days post-inoculation, we recorded the numbers of lightly and deeply colored leaves to estimate the strength of the immune response in the tobacco leaves. Three days after inoculation of *N. Benthamiana* plants with TSWV, we observed virus infection in all test groups. However, the percentage of light-red (indication of strong resistance) and dark-red (indication of weak resistance) leaves is quite different among the groups. In the Control group, 8 among the total 12 (∼67%) of the infected leaves turned dark-red color. However, only 2 or 3 among total 12 (∼21%) of the leaves showed dark color in SA-treated plants (Figure 8A). There are no major differences between SA-MN, SA-Spray, and SA- Infiltration groups. After 6 days, nearly 92% of the infected leaves were dark red in the Control group. In contrast, the percentage of dark-red leaves in SA-treated groups was kept lower at around 50% (Figure 8A).

**Figure 8.**
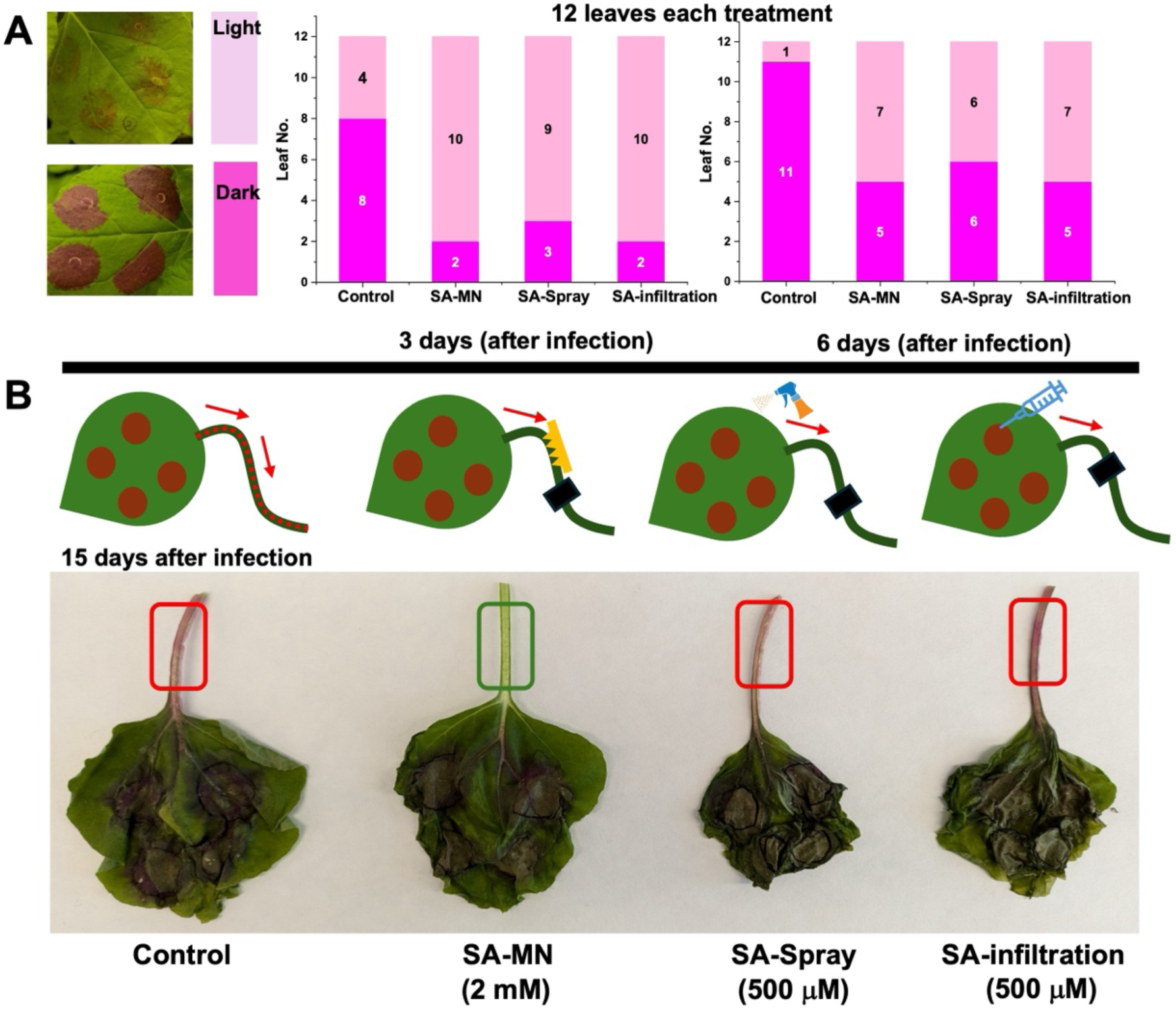
**SA-MN delivery prevents TSWV movement in *Nicotiana Benthamiana***. A) TSWV infection in *N. benthamiana* leaf after 3 days and 6 days of inoculation, respectively, as evident by the expression of RUBY. The light-red color indicated less TSWV infection, while the dark-red color indicated higher TSWV load and less pathogen resistance. B) The phenotype of TSWV infected-*Nicotiana benthamiana* leaves (15 days post infection). SA-MN delivery inhibited the TSWV movement from the leaf to the petiole.

This indicates MN is a highly effective delivery platform for sending SA into leaf tissues and inducing TSWV resistance. It is also interesting to note that after 15 days of viral infection, all infected leaves turned a dark red color and died due to virus infection. The petioles of Control, SA-Spray and SA-Infiltration groups all turned red color (Figure 8B), indicating the transport of TSWV along the plant vasculature system. However, the SA-MN treated petiole remained green (Figure 8B), indicating a strong viral resistance and fewer viral transport in the SA-MN treated petiole tissue.

The comparison experiments were repeated by increasing the concentration of SA in SA- Spray and SA-Infiltration groups to 1 mM instead of 500 μM (Figure S16A). After 20 days of infection, the Control group clearly showed that the movement of TSWV from the leaf tissue to the stem, along the vein in the WT *Nicotiana Benthamiana* plants (Figure S16B). However, in SA-MN, SA-Spray, and SA-Infiltrated plants, no red color on the stem was observed (Figure S16B). After 30 days of infection, the Control plant revealed a dwarf phenotype compared to the SA-treated plants (Figure S17A). The stem of Control plants showed tissue wilting. On the contrary, the SA-MN and SA-Spray treated plants were still healthy, while some viral infection (red color) was also found in SA-Infiltrated plants (Figure S17B), which may be due to less total SA treatment dose in the infiltration method (Infiltration: 5 mL of 1 mM SA; Spray: 10 mL of 1 mM SA and repeated 2x).

These experiments strongly suggested that SA could be effectively delivered to *Nicotiana Benthamiana* plants by the MN delivery method and induce resistance to TSWV infection. Compared to the traditional spray method, although the SA-MN method is more frequently applied (totally 4x) and has a higher SA concentration (2 mM), it significantly reduces the total application dose by ∼99% (Table 1) while achieving similar immunization effects. Moreover, the spray method required multiple treatment sites, including applications to the adaxial and abaxial sides of the leaf, while the PVA-MN injection simply works on the stem or petiole. This suggests that our PVA- MN delivery method not uses less active ingredient but also could be potentially applied faster due to easier application locations.

## Discussion

In this study, we demonstrated that PVA microneedle (MN) patches could be effectively utilized as a precision plant delivery system for various compounds, including small-molecule dyes, growth factors, and antiviral hormones (Figure 9). Compared to conventional delivery methods such as foliar spray and soil application, which suffer from high chemical losses, environmental impact, and low absorption (Boonupara *et al*., 2023; Sinha *et al*., 2022; Zeng *et al*., 2021), PVA- MN offer a promising alternative that can directly deliver target molecules into plant tissues with high precision and low application dose and environmental side effects.

**Figure 9.**
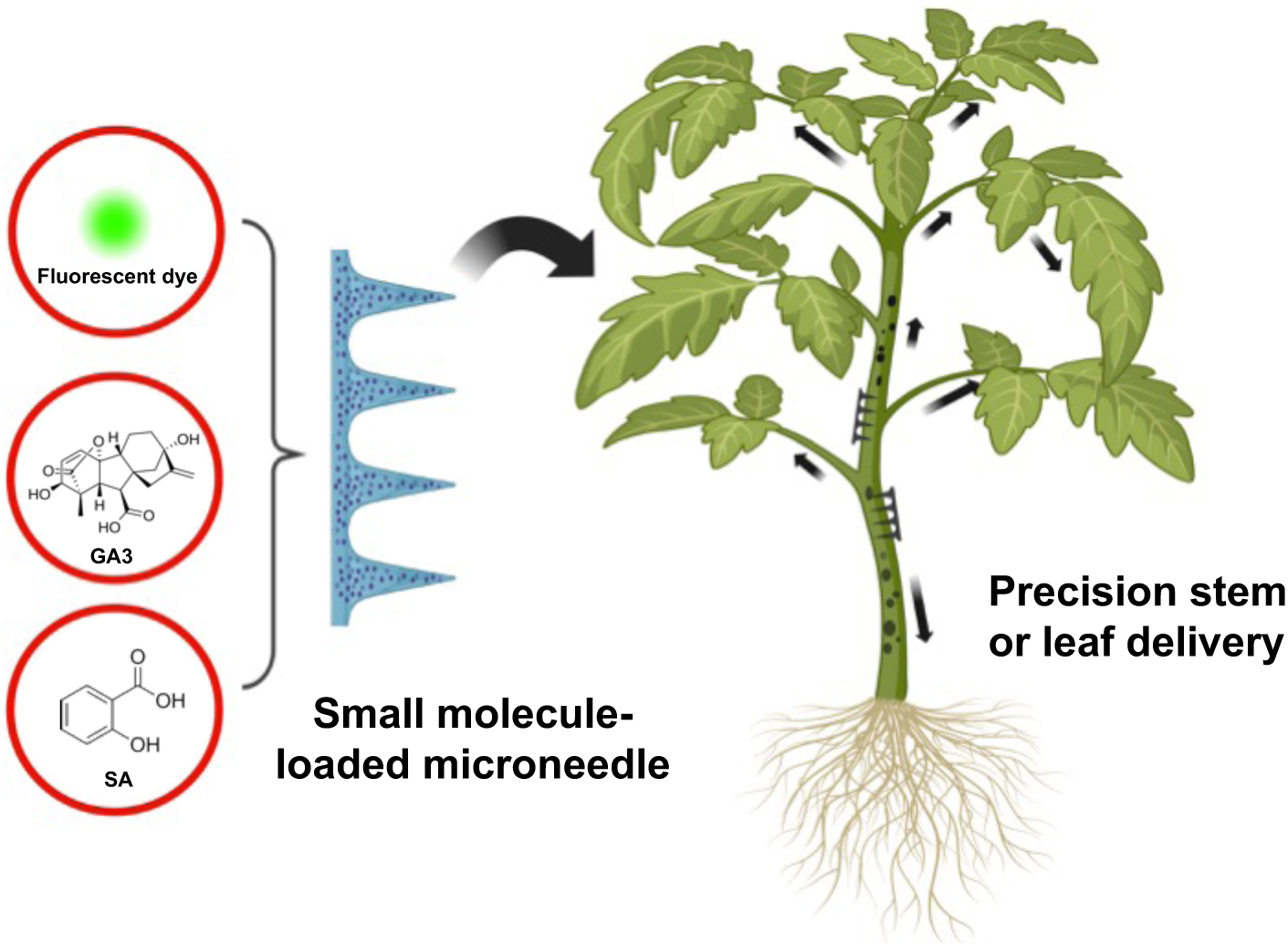
Summary of different cargos delivered by the PVA-MN platform into a live plant in this study.

Our results show that PVA MN successfully delivered fluorescent dyes into tomato stem vascular tissues, demonstrating their ability to penetrate plant tissues and promote compound diffusion (Figure 1). This highlights the advantage of PVA-MN in enhancing plant uptake and transport of bioactive molecules, which are often limited by environmental factors and plant surface barriers. Wounding assessments indicated that although MN insertion causes initial mechanical damage, the negative effects on stems are temporary and can be recovered by the plant itself (Figure S1-S5). Delivery of GA3 via PVA-MN accelerated tomato stem growth and altered chlorophyll levels in new leaves, confirming effective hormone delivery (Figure 2). Compared with foliar spraying, MNs produced comparable or slightly improved growth responses while reducing GA3 usage by nearly 90%, indicating a more controlled and efficient delivery method. Upregulation of GA receptor genes further confirmed successful activation of hormonal signaling. In Arabidopsis ft-10 mutants, MN-mediated GA3 delivery induced early bolting similar to foliar spraying, validating MN efficiency in a model system (Figure 4). The increased expression of GA receptor genes underscores the potential of PVA-MN for precise regulation of plant growth and development in agricultural applications. Finally, the application of SA via PVA-MN in *N. benthamiana* demonstrated the MN’s ability to enhance plant resistance against TSWV, a new way of plant immunization with high accuracy (Figure 8). The restricted viral movement in SA-treated plants highlights the effectiveness of MN in delivering antiviral compounds directly to the infection site, thereby enhancing the plant’s natural defense mechanisms. This approach offers a targeted method to combat viral infections, potentially reducing the need for broad-spectrum chemical treatments that can harm the environment.

Our PVA-MNs differ from several previously reported microneedle systems used in agriculture. For example, Cao and colleagues designed a silk-based microneedle platform (Cao *et al*., 2023; Cao *et al*., 2020) that can also deliver small molecules and Agrobacterium to leaf tissues. However, a major limitation of their approach is that the fabrication process for silk microneedles is more complex and the production cost is much higher than that of our PVA-MNs.

On the other side, while several emerging nanomaterial-based delivery methods have been demonstrated for delivering small molecules as well as DNA fragments and even proteins (Demirer et al., 2021) to plants, these approaches are more suitable for small-scale laboratory studies. Their overall cost is higher than that of PVA MNs, and their scalability is comparatively more limited (Beig *et al*., 2022; Demirer et al., 2019; Demirer et al., 2020). In addition, while nanomaterials may enhance delivery at the cellular level, at the tissue level, many nanomaterials still rely on foliar spray or infiltration-based delivery approaches. As the initial demonstration of using PVA MNs for molecular delivery in plants, we only selected several small molecules such as plant hormones for validation in this study. In the near future, MN also holds great promise for the delivery of large biomolecules—including DNA, RNA, and proteins, even the gene editing systems—into plant tissues.

Compared with existing or other emerging plant delivery methods, the MN approach demonstrates obvious advantages. Owing to their simple fabrication, low cost, and environmental safety, MN patches can be used for direct attachment to plant stems, petioles, or fruit pedicels for localized delivery of nutrients, hormones, or protective agents. In greenhouse settings, MNs can be deployed manually or via automated dispensers to achieve controlled and repeatable dosing. In field applications, MN systems could be integrated with existing agricultural machinery systems for large-scale, site-specific treatment. Furthermore, the smart polymer matrix allows the design of time-controlled or humidity-responsive release formulations, enabling sustained molecular delivery under fluctuating environmental conditions. After use, PVA MNs can be dissolved in water or degraded by various microorganisms under aerobic and anaerobic conditions, producing CO₂ and H₂O rather than persistent plastic fragments, therefore minimizing microplastic contamination in plant tissues or the surrounding environment (Malka and Margel, 2023; Nair and Laurencin, 2007; Tokiwa and Calabia, 2007). Together, these features demonstrate the feasibility of applying MN technology in precision agriculture to enhance crop health and productivity while minimizing environmental impacts.

Despite these promising results, several challenges remain in implementing MN-based delivery methods in future. For instance, the dehydration effect observed in MN-treated leaf tissues suggests that MN application may be more suitable for stems or stronger plant tissues. Stem delivery may potentially also accept a larger amount of payloads and transport the delivered payloads faster to other parts of the plant than leaf delivery. Moreover, due to the hydrophilic properties of PVA, it is essential to maintain the stability and activity of loaded biomolecules within the PVA matrix, which requires further optimization of the microneedle formulation.

In the future, the integration of PVAMN technology with agricultural robotics offers another direction for next-generation precision crop management. As field robots and autonomous platforms continue to advance in sensing, navigation, and manipulation (Oliveira et al., 2021; Vougioukas, 2019; Wakchaure et al., 2023), MN-based delivery systems could be incorporated into robotic arms or end-effectors to enable fully automated, site-specific treatment of individual plants. Such systems could autonomously diagnose nutrient deficiencies, pathogen infections, or growth abnormalities using onboard imaging and sensor networks, followed by immediate, localized intervention through MN-mediated delivery with minimal human involvement. The soft and adaptable nature of PVA-MN patches also makes them compatible with gentle robotic grippers, reducing the risk of mechanical damage during automated application. In large-scale production systems, fleets of robotic units could deploy MN patches with high precision and consistency, allowing repeated or need-based delivery that responds dynamically to plant developmental stages or environmental fluctuations. This convergence of MN-based biochemical delivery and intelligent robotic actuation holds the potential to create a highly responsive, data-driven agricultural framework that maximizes resource efficiency while minimizing chemical and labor input.

In summary, our findings suggest that PVA-MNs are a viable and effective tool for plant compound delivery, offering benefits over traditional methods in terms of precision, efficiency, and reduced environmental impact. Future research should focus on scaling up MN fabrication processes, automating delivery techniques, and expanding the range of compounds that can be effectively delivered through this innovative system. With continued development, MN delivery technology could revolutionize precision agriculture, enabling more sustainable and targeted interventions in crop management.

## Materials and methods

### Plant materials and growing conditions

Arabidopsis thaliana *ft-10* mutant seeds (TAIR Germplasm CS9869) and ecotype Columbia (Col- 0) seeds were sterilized by soaking for 1 min in 70% ethanol and then for 10 min in 20% bleach, followed by five to seven rinses with sterile distilled water. Sterilized seeds were placed on Murashige and Skoog (MS) plates and kept at 4 °C for 2 days. The plate was placed in a plant growth chamber (Caron 7304-22-1) at 22 °C with long day (day/night 16h/8h) conditions, 70–100% relative humidity, and light intensity of 100 µmol m^−2^ s^-1^. Seedlings were transferred to pots (2.5 × 2.5 × 3 in^3^) with soil (Miracle-Gro Moisture Control Potting Soil Mix) on day 7 and watered regularly. Tomato (*Solanum lycopersicum* var. Beefsteak) seeds were purchased from The Seed Plant. Tomato seeds were grown in pots (3.5 × 3.5 × 3 in^3^) in the plant room under long day (day/night 16h/8h) at 80 µmol m^−2^ s^−1^, 70–90% relative humidity, day and night temperatures of 26 and 20 °C, respectively. *Nicotiana benthamiana* plants were grown in the growth chamber under 12h day and 12h night cycles at 26°C and 275 µmol m^−2^ s^−1^.

### TSWV virus infection

*Agrobacterium tumefaciens* GV3101 carrying recombinant plasmids (pL, pM-RUBY, pS-NSs, and triple VSR (viral suppressor of RNAi) were grown in Luria Broth media containing appropriate antibiotics (Rifampicin (50 μg/ml), Kanamycin (50 μg/ml), and Gentamycin (50 μg/ml)) for overnight. Cells were harvested from overnight grown cultures of *Agrobacterium* and re-suspended into the infiltration buffer (10 mM MgCl2, 10 mM MES pH 5.6 and 100 μM acetosyringone) at OD600 of 1 and further incubated at 28^ο^C for 3 hours. Before infiltration, *Agrobacterium* cultures carrying pL, pM-RUBY, pS-NSs, and triple VSR plasmids were mixed at final OD600 of 0.25. Three hours post-incubation, fully expanded *Nicotiana benthamiana* leaves were co-infiltrated with *Agrobacterium* cells harboring pL, pM-RUBY, pS-NSs, and triple VSR plasmids at final OD600 of 0.25 (Liu *et al*., 2023).

### PVA MN fabrication process

The microneedle patch was fabricated using a simple centrifugation method using polydimethylsiloxanes (PDMS)-based negative molds (Blueacre Technology) in a 10 (w/v) % polyvinyl alcohol (PVA) (M.W. 31,000-50,000, 98.0-98.8% hydrolyzed) (Acros Organics) solution. To remove dust and debris from the micron-scale crevices of the PDMS molds, each chip is first cleaned by repeated rinses with deionized (DI) water followed by two rounds of sonication for 5 minutes each. The molds then dried at 35°C for 10-20 minutes to evaporate any remaining water droplets clinging onto the needle edges. Next, 1.2 mL of PVA is allotted in 1.7 mL centrifuge tubes, which undergo their first round of centrifugation at 6260g (Fisher Scientific accuSpin Micro17) for 10 minutes remove any trapped air bubbles. The molds are then submerged in degassed PVA solutions with the needle molds facing inwards and centrifuged again at 6260g for 10 minutes to fill the molds with PVA. The molds are then removed and sealed with another layer of PVA to form a strong base for the needles upon drying. Lastly, the MN patches are partially covered (for slow evaporation) and cured for 24 hours at room temperature to allow complete solidification.

*MN patch fabrication with Gibberellic acid (GA3):* 34.6 mg of GA3 is dissolved in 500 µL of dimethylsulfoxide (DMSO) and 500 µL of deionized (DI) water to make a stock solution of 100 mM of GA3. 6 µL of the solution is added to 1.2 mL of 10 (w/v) % PVA solution to make a final GA3 concentration of 500 µM in 10 (w/v) % PVA solution for one microneedle patch. Each 20 x 20 MN array is made from 500 μL PVA-GA3 solution. In the tomato treatment experiments, only a 4 x 10 array was pressed to the plant stem due to the small size of plant stems. The estimated amounts of total GA3 delivered to different plants by MN are summarized in Table 1.

*MN patch fabrication with Salicylic Acid (SA):* 11.04 mg of SA is dissolved in 15 mL of DI water to make a stock solution of 5.33 mM. This solution is then added to 16 (w/v) % PVA solution to make a final solution of 2 mM SA and 10% (w/v) PVA for microneedle fabrication.

*MN patch fabrication with Rhodamine and 5(6)-carboxyfluorescein diacetate (CFDA):* 52 mg of Rhodamine is dissolved in 1.2 mL DI water to make a stock solution of 0.09 M. On the other hand, to make a stock solution of 0.27 M CFDA, 25 mg of the powder is dissolved in 200 µL of DMSO. 33 µL and 11 µL of the respective stock solutions were added to 30 mL of 10 (w/v) % PVA to make a final solution containing 0.1 mM Rhodamine and CFDA, respectively, which was used to fabricate the microneedles.

### Fluorescent dye delivery in tomato plants

CFDA (1 mM) and rhodamine 6G (1 mM) were mixed with a DMSO solution at a volume ratio of 1:100 when fabricating the PVA-MN. To monitor fluorescence signal from the delivery and transport of CFDA and rhodamine 6G, cross-sections upstream and downstream along the petiole were observed under a microscope with fixed light intensity and an exposure time of 20 ms. The petiole was also sliced longitudinally to image the distribution profile of CFDA in the xylem. Image analysis was performed using ImageJ 1.52i. Fluorescence intensity was used as a proxy for the concentration of CFDA within the tested range, and fluorescence signals were integrated radially.

### Plant hormone preparation

GA3 Solution Preparation: A 0.1 M stock solution of GA3 was prepared by dissolving 34.6 mg GA3 in 1 mL 75% ethanol solution. GA3 spray solution was prepared by diluting the stock solution 1000× to a final concentration of 100 μM. Silwet-77 was added to obtain a final concentration of 0.01% in the GA3 spray.

SA Solution Preparation: A 100 mM stock solution of SA was prepared by dissolving 138 mg SA in 1 mL 75% ethanol solution. The SA spray solution was prepared by diluting 100 mM SA stock 1000× to the final concentration of 0.1x10^-3^ M. Tween-20 was added to obtain a final concentration of 0.01% in the SA spray and used as a surfactant to promote SA absorption.

### RT-qPCR based gene expression analysis

RT-qPCR analysis was performed using SYBR-Green PCR Master mix (Invitrogen, Carlsbad, CA, United States) on a CFX96TM (Bio-Rad, California, CA, United States). Gene-specific primer pairs and different thermal programs (Supplementary Table S1) were developed to compare expression levels in *Arabidopsis*, tomato, and *N. benthamiana* plants. *Arabidopsis UBQ10* gene and tomato *Actin* gene were used as reference controls for normalization. A pre-RT-qPCR analysis showed that the reference gene expression is stable under the plant hormone treatments including GA3. Amplified products were monitored using an optical reaction module (SYBR-green) and the resulting amplified values of genes were normalized against those of references according to a previous protocol for plants (Li et al., 2021). The steps of qPCR analysis were carried out following the method described previously (Li *et al*., 2021). The reaction system was composed of 12 µL mixture, including: 10 µL of SYBR-green-preMix, 1.0 µL of 10 µM forward and reverse primer, and 1.0 µL of cDNA template (Li et al., 2017). The thermal program followed the protocol of Bio-rad SYBR-Green PCR Master-mix. The thermal program is 95 °C for 30s, (95 °C for 10s, 60 °C for 1 min) ×40, 65 °C to 95 °C with 0.5 °C increments in 2s, 4 °C. Three replicates were performed for each gene expression analysis. All the primers were listed in Supplementary Table S1.

### Salicylic acid (SA) extraction from tobacco leaf and detection using the HPLC-MS/MS

For SA analysis, approximately 100 mg of fresh leaf tissue is collected and ground into a fine powder in liquid nitrogen. The powder is transferred into a 2-mL microcentrifuge tube and extracted with 1.0 mL of extraction solvent (methanol:water:formic acid = 80:19:1, v/v/v), followed by the addition of a known amount of internal standard. The mixture is vortexed for 1 min and sonicated in an ice-cooled ultrasonic bath for 10–15 min, then centrifuged at 15,000 g for 10 min. The supernatant is transferred to a new tube. The pellet is re-extracted with an additional 0.5 mL of extraction solvent, and the supernatants from both extractions are combined (total volume ∼1.5 mL). The extract was stored at -80°C before measurement or taken directly to SPE purification if preferred.

The content of SA was measured on an HPLC–MS/MS System (SCIEX Triple Quad 5500) equipped with a C18 column (Phenomenex, 4.6 × 250 mm, 5 μm). The injection volume was 5 μL with a flow rate of 0.4 mL/min. The column pressure is 1800 Bar and a column temperature of 40°C. The flow channel is A:100% water and B:100% Acetonitrile. A total 12 min timetable is: Start at 95% A, 5% B; 1 min: 95% A, 5% B; 6 min: 5% A, 95% B; 7.5 min: 5% A, 95% B; 10 min: 95% A, 5% B; 12 min: 95% A, 5% B. The mass spectrometer was operated under negative ionic mode with the Electrospray Ionization (ESI). The gas temperature was set to 320°C. The gas flow rate was set to 6 L/min, and the temperature of sheath gas, with 11 L/min flow rate, was set to 350°C. The data were acquired and analyzed with Analyst software. The SA were identified by comparing the standard substances. In result analysis, the characteristic ion fragments of the SA were set as 137/93 or 137/65. We observed that the SA retention time is at 7.29 min, and the 137/93 peaks were the major primary and secondary ion fragments in our analysis method.

## Author Contribution

M.L. contributed to experimental design, plant treatments, RNA extraction, analysis of all experimental data, figure preparation, and manuscript drafting. A.D.P. contributed to PVA-MN preparation. J.X. contributed the SEM characterization of PVA-MN. H.J. contributed to RNA extraction from the plant tissue and RT-qPCR analysis, and the manuscript editing. Z.Z. contributed to the tomato plant cultivation, GA3, SA treatment, and the SA extraction from the tomato plants. Y.H. and N.L. contributed the HPLC- MS/MS analysis of SA from plants. Y.L., L.G., and T.X. provided the HPLC- MS/MS platform and contributed to the SA concentration analysis in the plant. D. S. and A.E.W. helped construct the RUBY-TSWV and TSWV infection experiments. Q.W. perceived and designed the project, developed experimental plans, supervised data analysis and figure preparation. All authors contributed to the revision of the manuscript.

## Acknowledgements

The authors thank Dr. Wusheng Liu at NC State Department of Horticulture for helping with RNA extraction. We also thank Lydia Parks for developing the MN fabrication protocol.

