## Supporting information for "Microneedle-based precision payload delivery in plants"

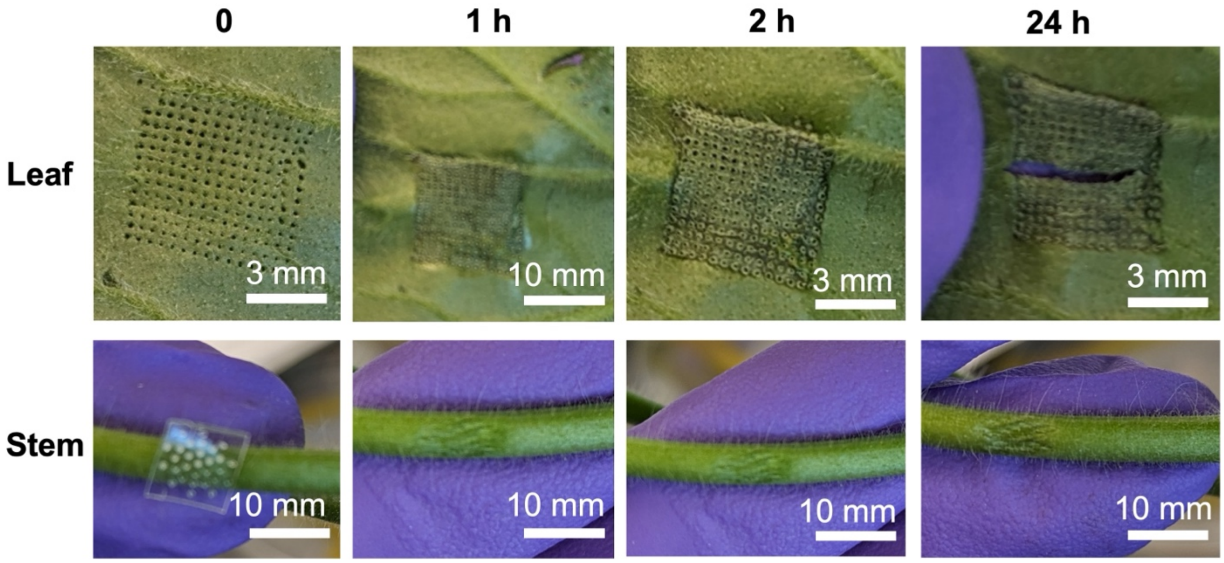


**Fig. S1: The effect of PVA MN punch on the plant leaf and stem.** The punch position at leaf started to dehydrate after 1 hour (top row, MN contact time: 30 s). Severe dehydration and tissue cracking were observed after 24 hours of MN punch (top row). In contrast, the stem is more robust against the MN punch (bottom row).

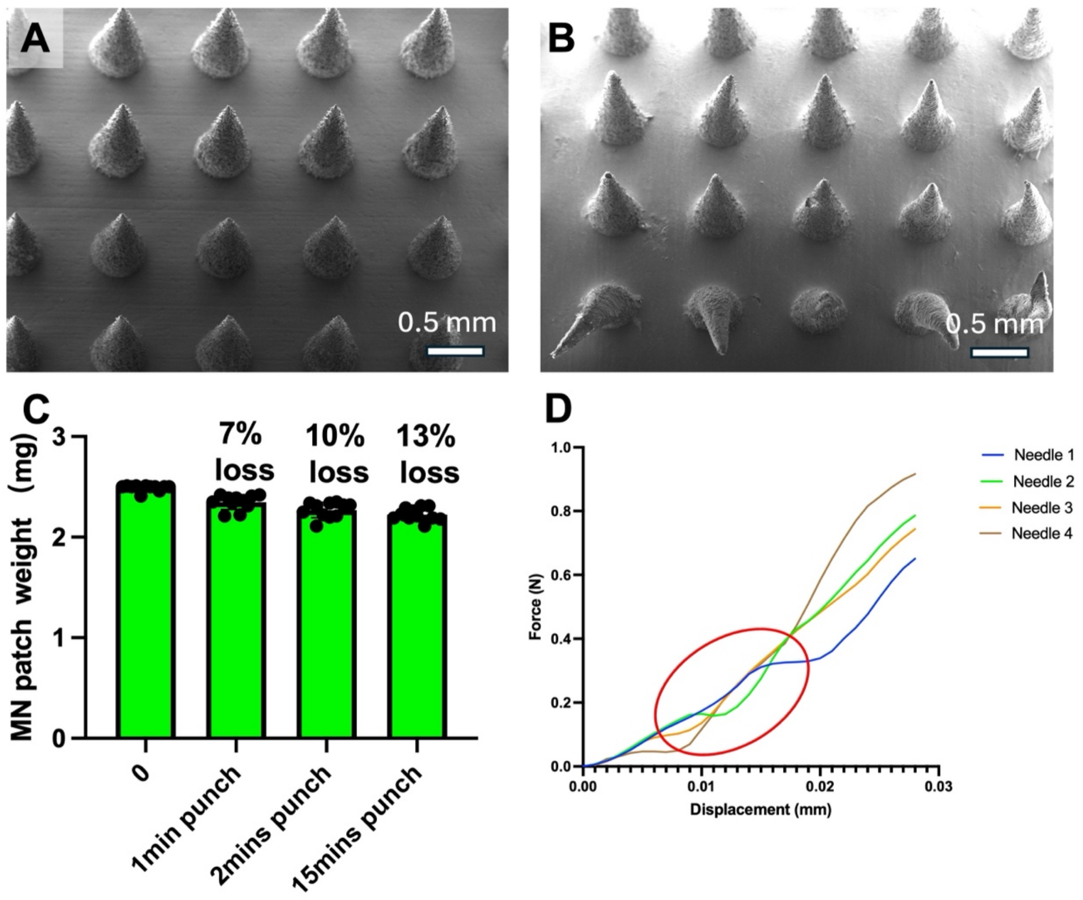


**Fig. S2: Characterization of MN punching on the tomato stem.** A) Representative SEM image of PVA-MN patch before stem punching. B) Representative SEM image of MN patch after stem punching (contact time: 30 s). C) The average weight of the PVA-MN before and after injection on the tomato stem for different contact times of 1, 2, and 15 min (n=10). D) Mechanical behaviors of PVA-MN when penetrating the tomato stem under compression (n = 4 repeats). The red circle indicates the needle fracture points during leaf tissue penetration.

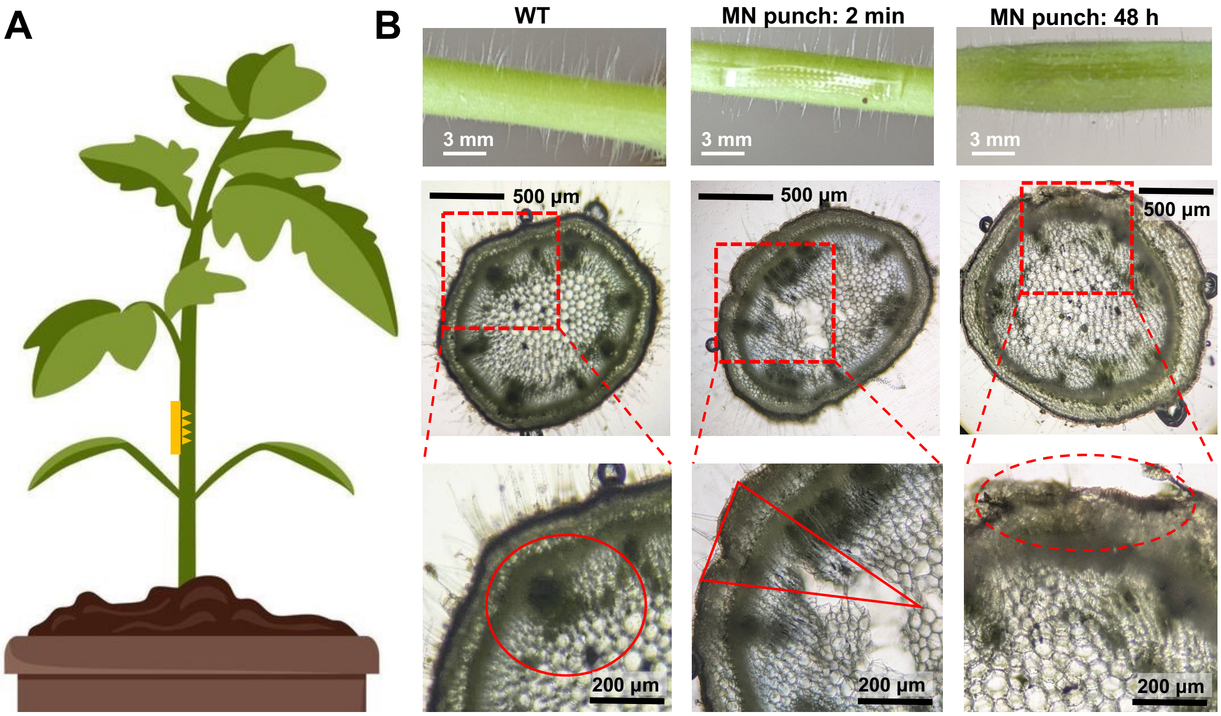


**Fig. S3: The cross-section of the tomato stem and wounding response to PVA MN punch.** A) Schematic of PVA-MN punch on the tomato stem. B) Photos and cross-section microscopy images of the WT tomato stem (without MN punch), 2 min, and 48 h after MN punch. The upper row is the photographs of the stem; the middle row is the cross-section images captured by a brightfield microscope with a 4X objective lens, and the bottom row is the 10X objective lens images. The red circle in WT indicates the vascular system in the normal tomato stem; The red triangle shows the microneedle injection site on the tomato stem; The red dashed circle in MN-punch-48h indicates the microneedle injection site on the surface of the tomato stem.

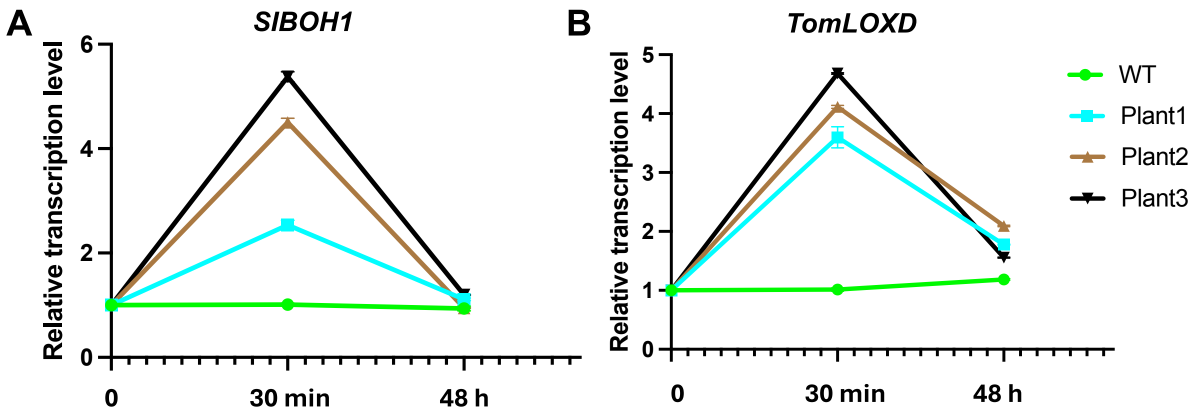


**Fig. S4: Gene expression of *SlBOH1* and *TomLOXD* after 30 min and 48 h of PVA-MN treatment.** A) The relative expression level of *SlBOH1* in tomato stem tissue as a function of time after PVA-MN punch. B) The relative expression level of *TomLOXD* in tomato stem tissue as a function of time after PVA-MN punch. n=4 (each plant has four RT-qPCR replicates).

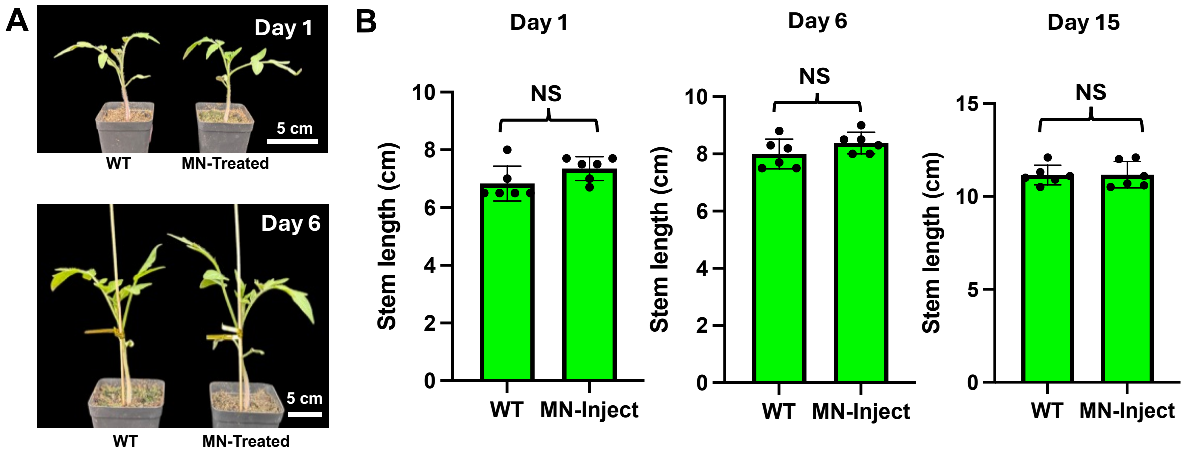


**Fig. S5: Effect of PVA-MN wounding (x3) on tomato growth.** A) The phenotypic observation of the WT and MN-punched tomato plants at Day 1 and Day 6. B) The stem length measurement results demonstrated that there are no obvious differences between the WT and MN-punched tomato plants from Day 1 to Day 15 (n=6).

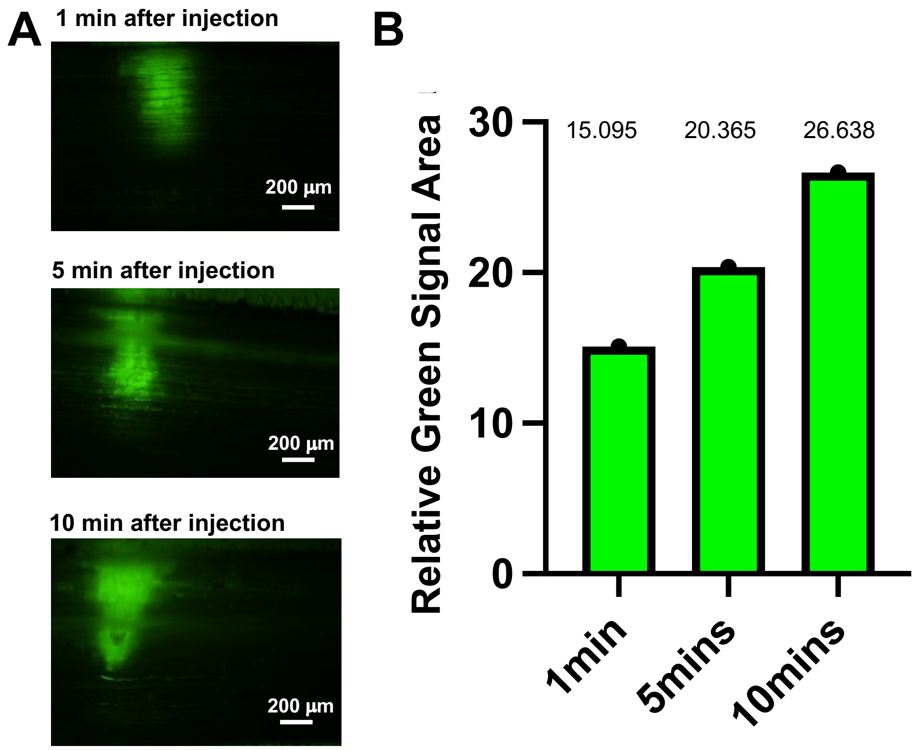


**Fig S6. Diffusion of MN-injected 5(6)-carboxyfluorescein diacetate (CFDA) fluorescent dye in tomato stem.** A) Cross-sectional fluorescence microscopy images showing CFDA delivered into stem tissue and transported along the xylem. B) The relative green signal area shows the diffusion of CFDA in the tomato stem after 1, 5, and 10 mins of injection.

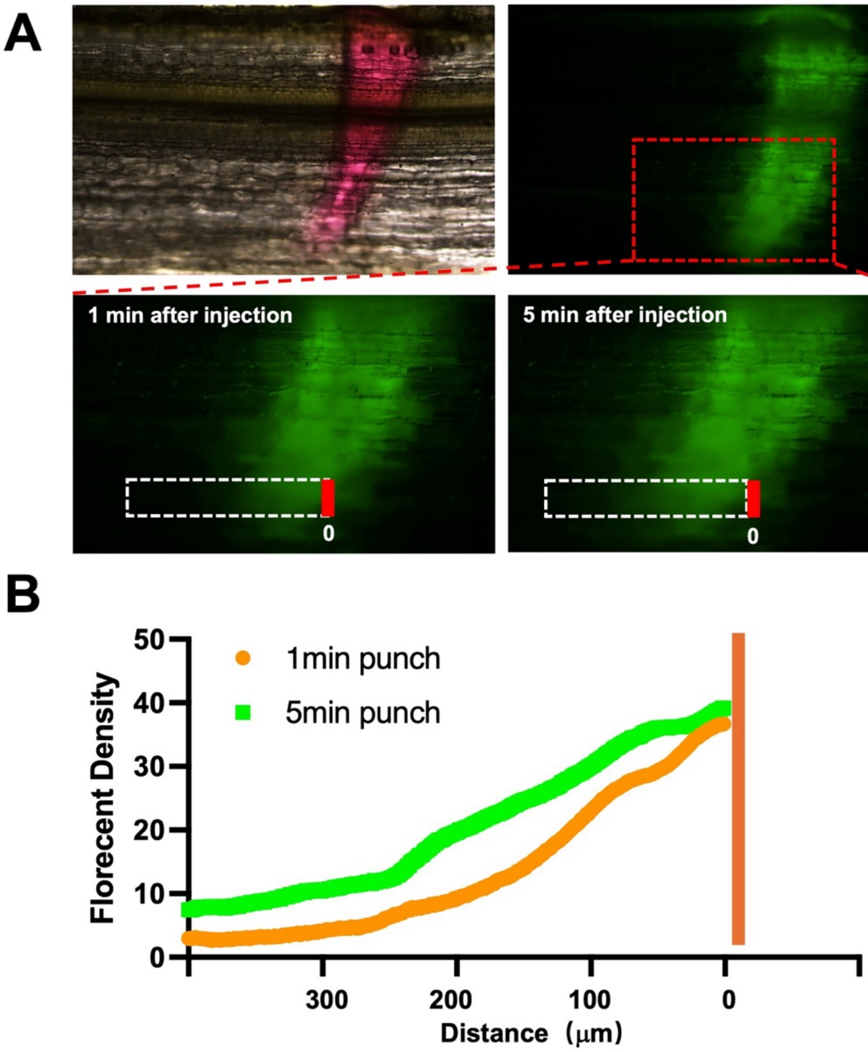


**Fig. S7: Movement of 5(6)-carboxyfluorescein diacetate (CFDA) fluorescent dye along xylem in tomato stem.** A) Brightfield and fluorescence microscopic images of CFDA distribution along the xylem. B) Corresponding fluorescent intensity analysis, depicting CFDA distribution along xylem (1 and 5 min postinjection, respectively). Red and green lines are the fluorescent density at 1-min and 5-min post-injection, respectively.

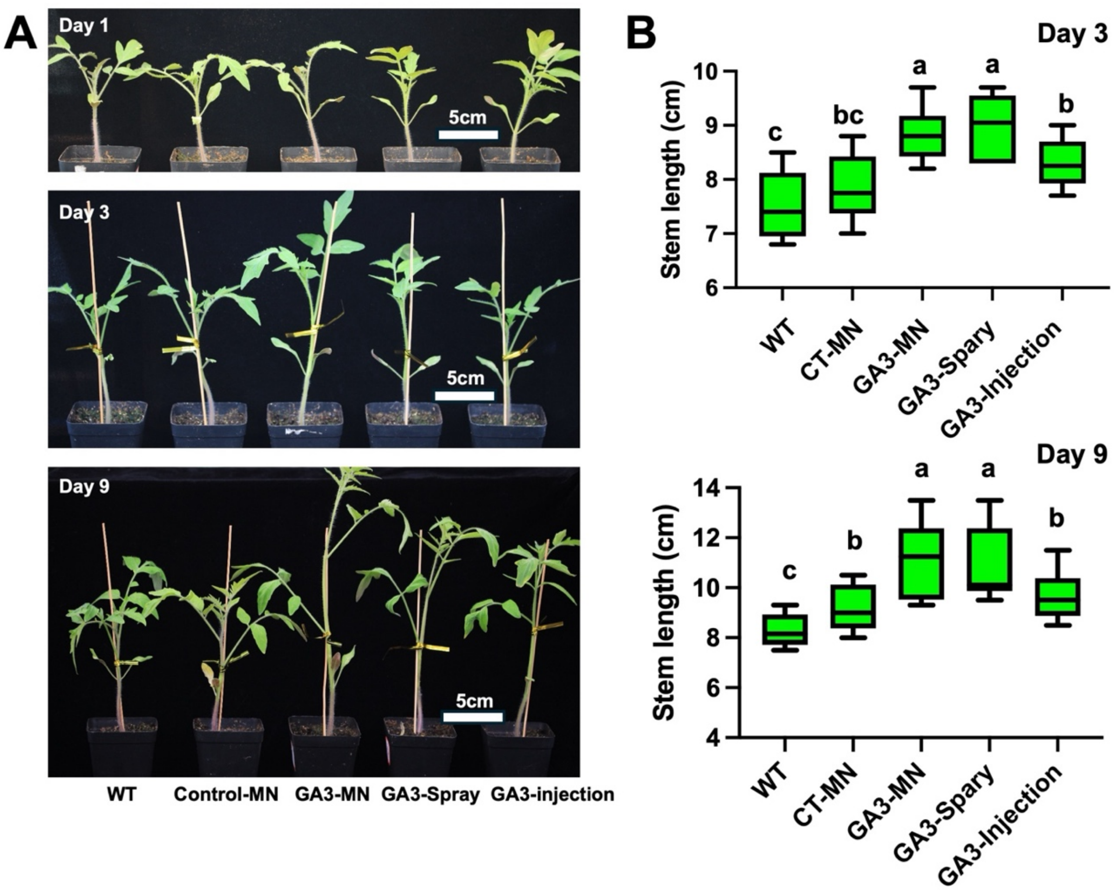


**Fig. S8: MN delivers GA3 into tomatoes and promotes stem growth.** A) After GA3 applications, the tomato stem showed different growth rates for different GA3 treatments. B) The stem length of the tomato plants at Day 3 and Day 9 under different GA3 stimulations (n=12). Different letters above the bars indicate statistically significant differences between groups ($P < 0.05$). Groups that share a letter are not significantly different ($P > 0.05$). CT = control.

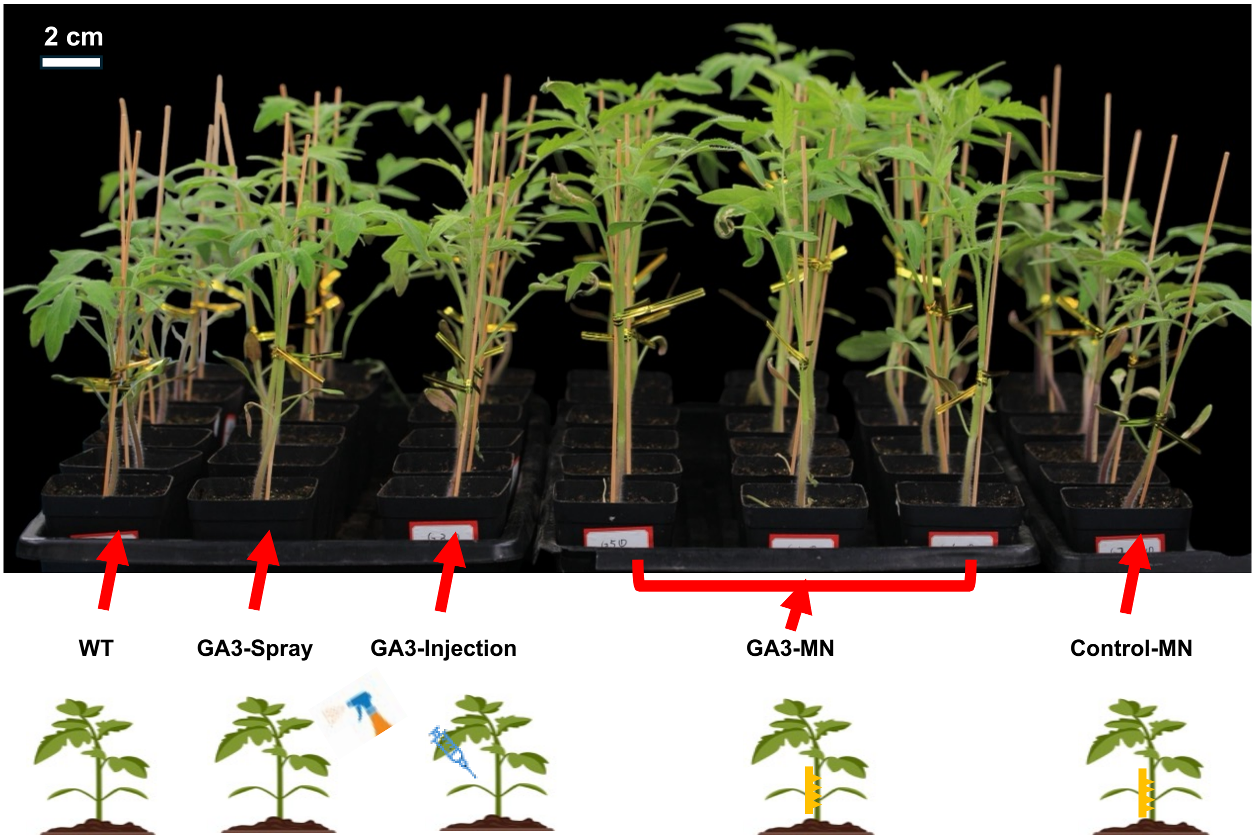


**Fig. S9:** An overview of all experimental tomato plants (including GA3-treated and non-treated plants) at Day 15 after treatment. The GA3-MN group outgrows all other controls.

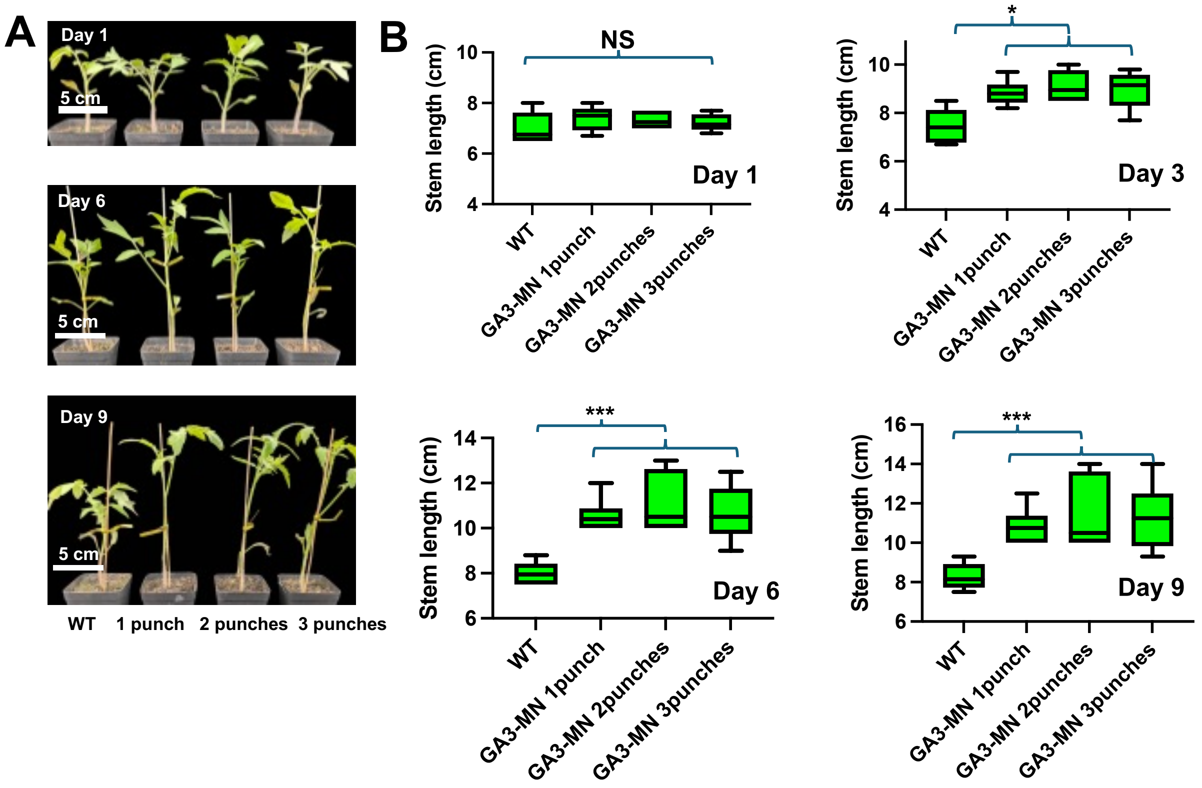


**Fig. S10: Effect of different numbers of GA3-MN treatment on tomato growth.** A) Phenotypic observation of tomato plants treated by 1x, 2x, or 3x GA3-MN at Day 1, Day 6, and Day 9, respectively. B) The stem length of the tomato plants at Day 1, Day 3, Day 6, and Day 9 under the different times of GA3-MN stimulations (n=12). (NS = not significant, $^{*}P < 0.05$, $^{**}P < 0.01$, $^{***}P < 0.001$)

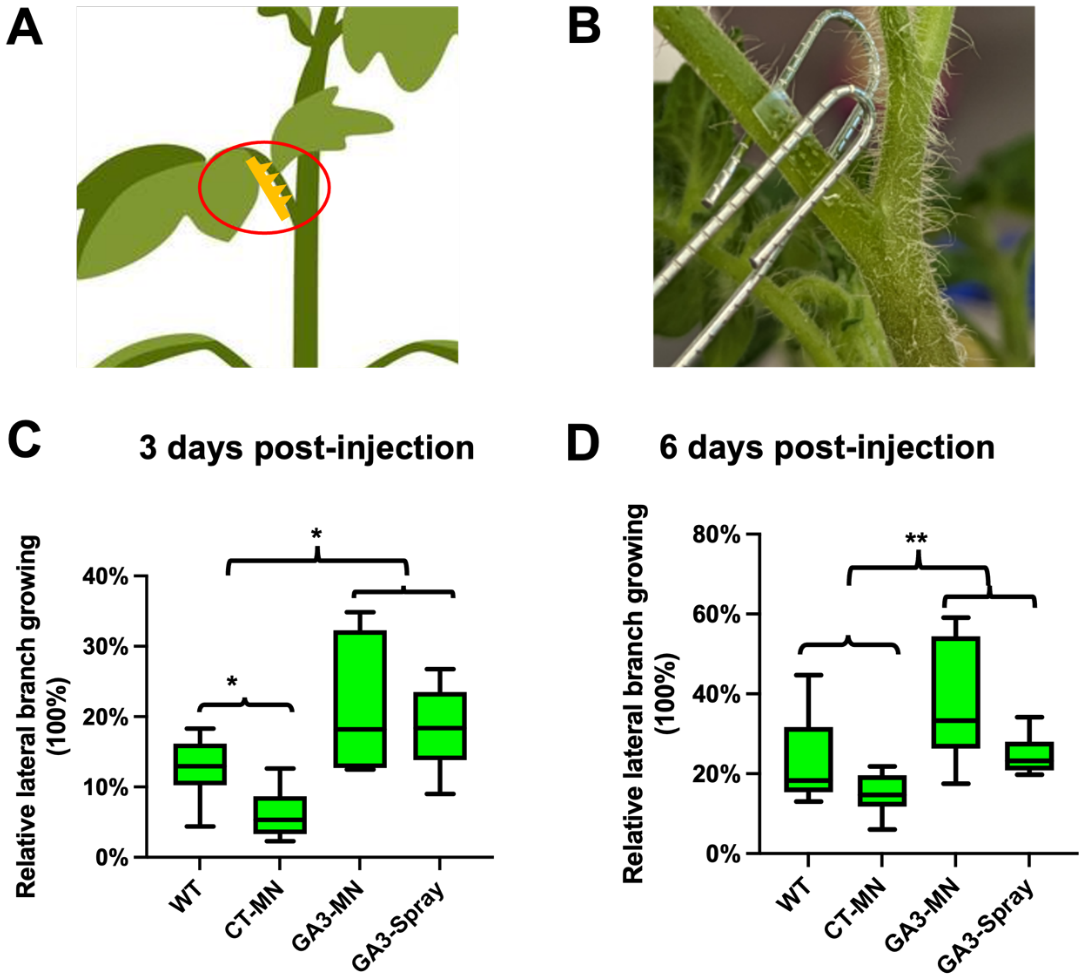


**Fig. S11: MN-GA3 treatment promotes petiole elongation in tomato plants.** A,B) Illustration and actual photo of GA3-loaded PVA-MN, punched on the lateral branch of the tomato plant. C,D) The relative lateral stem increases after different GA3 treatments at 3 days and 6 days (n=12). (NS = not significant, *P < 0.05, **P < 0.01, ***P < 0.001)

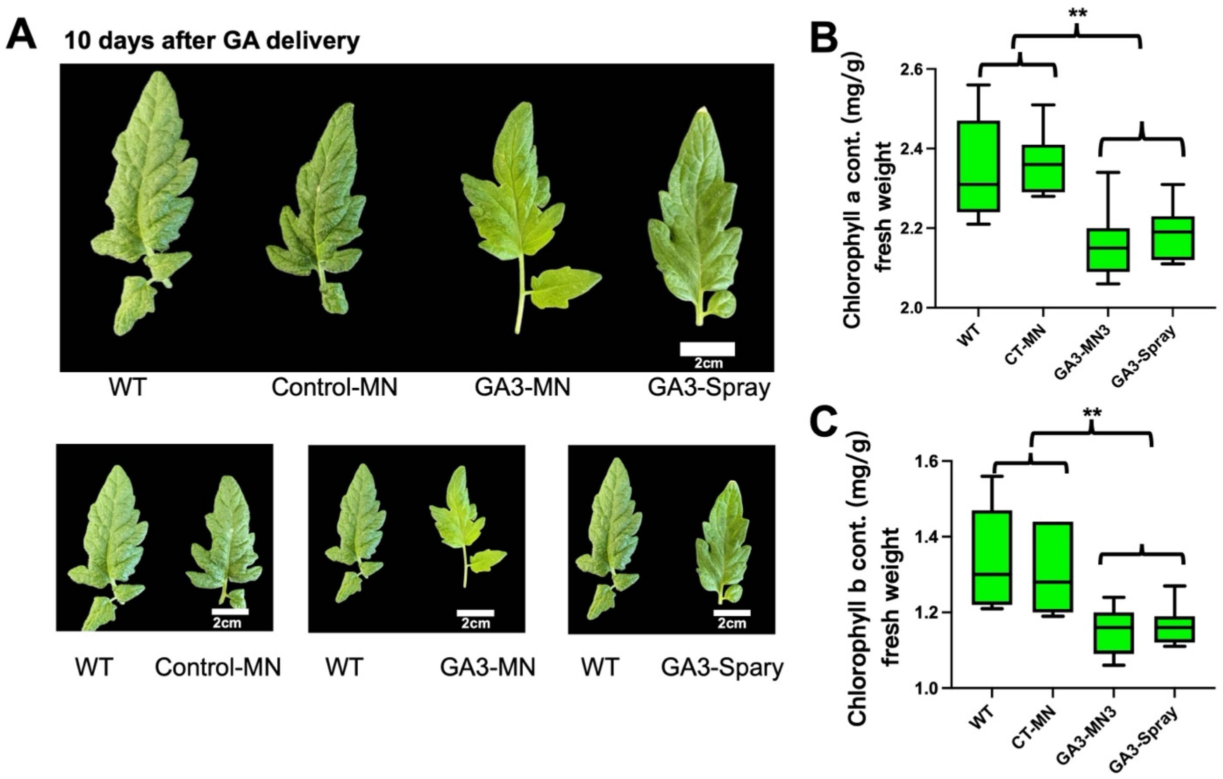


**Fig. S12: MN-GA3 treatment changed leaf color and Chlorophyll content.** A) The tomato plants in GA3-treated groups (GA3-MN and GA3-Spray) showed a noticeable color difference from 4th to 6th compound leaf (newly emerging leaves). B) The concentration of chlorophyll a ($p = 0.0049$) in the 4th to 6th leaves (mixed samples) (n=10). (NS = not significant, *$P < 0.05$, **$P < 0.01$, ***$P < 0.001$). C) The concentration of chlorophyll b ($p = 0.0025$) in the 4th to 6th leaves (mixed samples) (n=10). (NS = not significant, *$P < 0.05$, **$P < 0.01$, ***$P < 0.001$)

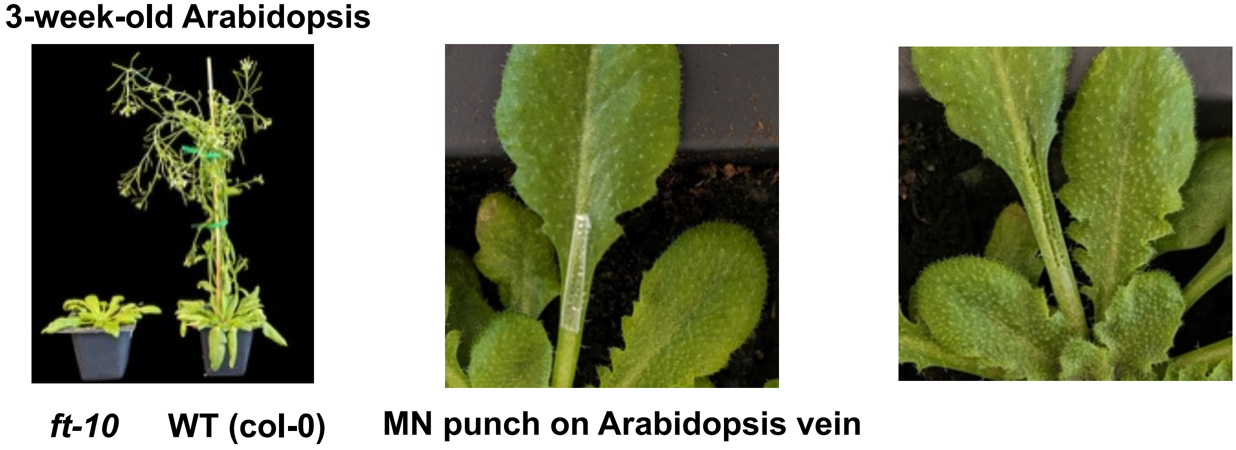


**Fig. S13: Lateral flowering phenotype of the *Arabidopsis* ft-10 mutant and wounding response of MN punch on the *Arabidopsis* plant.** A) Phenotype of the 21-day-old Arabidopsis ft-10 mutant and the WT (col-0). The WT plant flowers, but not the ft-10 mutant. B) The PVA-MN punch on the vein of *Arabidopsis*. C) The wounding response of the *Arabidopsis* vein after the PVA-MN injection.

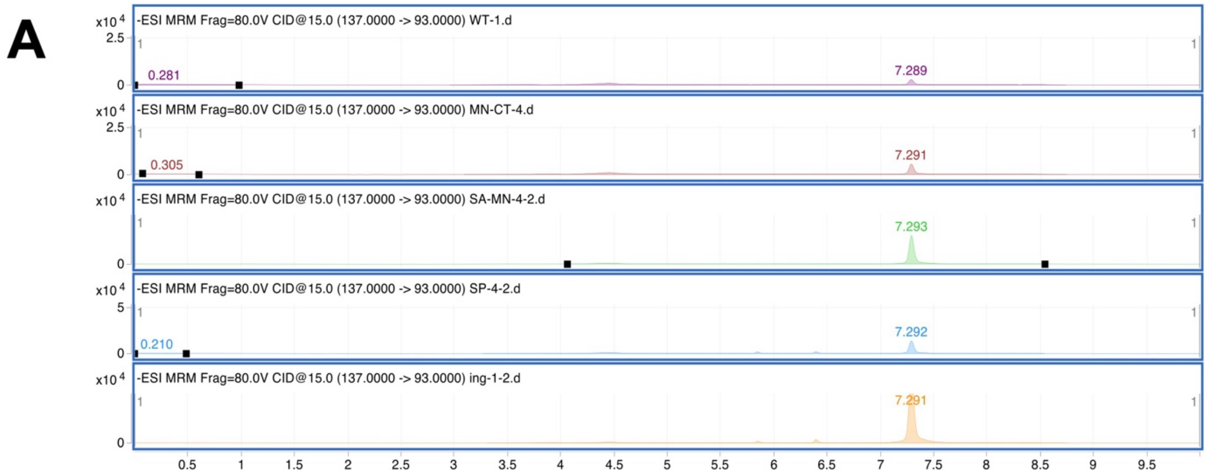


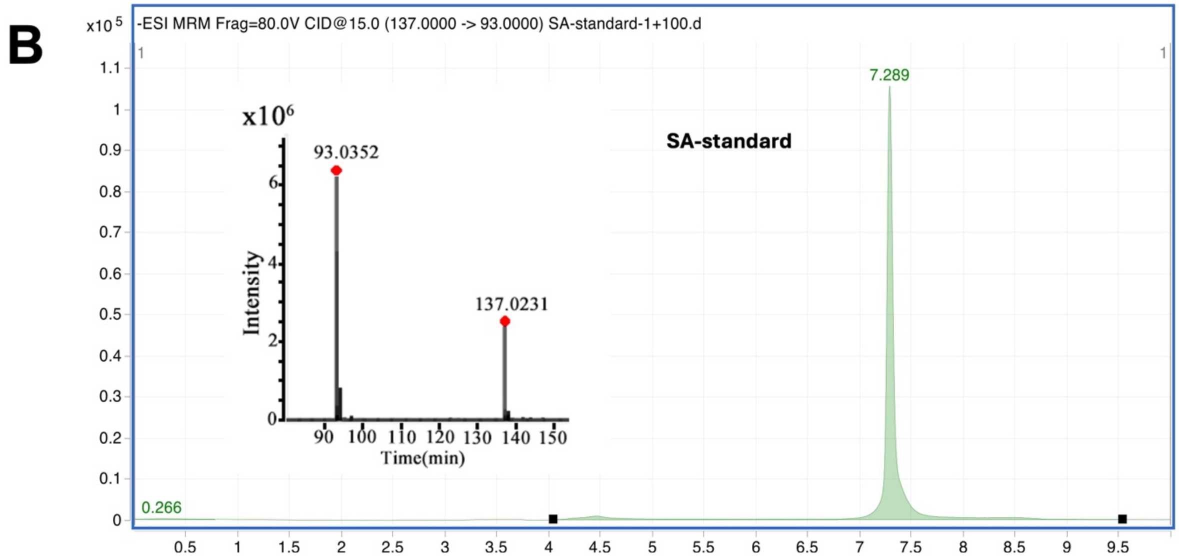


**Fig. S14: Plant tissue SA concentration analysis by HPLC-MS/MS.** A) HPLC-MS/MS analysis showed that the SA was successfully delivered into the leaf by using PVA-MN. The peak at 7.29 min is SA-MRM extraction through the HPLC-MS/MS analysis. B) HPLC-MS/MS spectrum of the SA standard. The 137 m/z and 93 m/z peaks represent the primary and secondary fragments of SA under the HPLC-mS/MS detection mode. The peak at 7.29 min is SA-MRM extraction through the HPLC-MS/MS analysis.

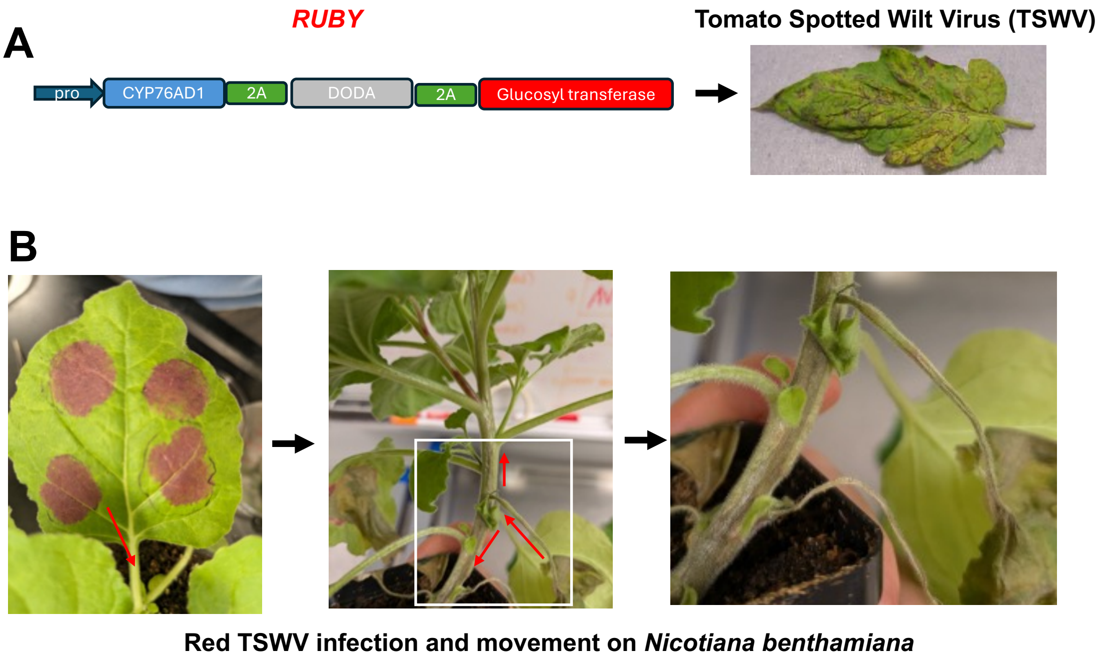


**Fig. S15: Infection of *Nicotiana benthamiana* with TSWV containing a Ruby reporter gene.** A) Illustration of recombinant tomato spotted wilt virus (TSWV) modified to express the RUBY reporter gene. B) The red phenotype after TSWV infection on *N. benthamiana* and its movement.

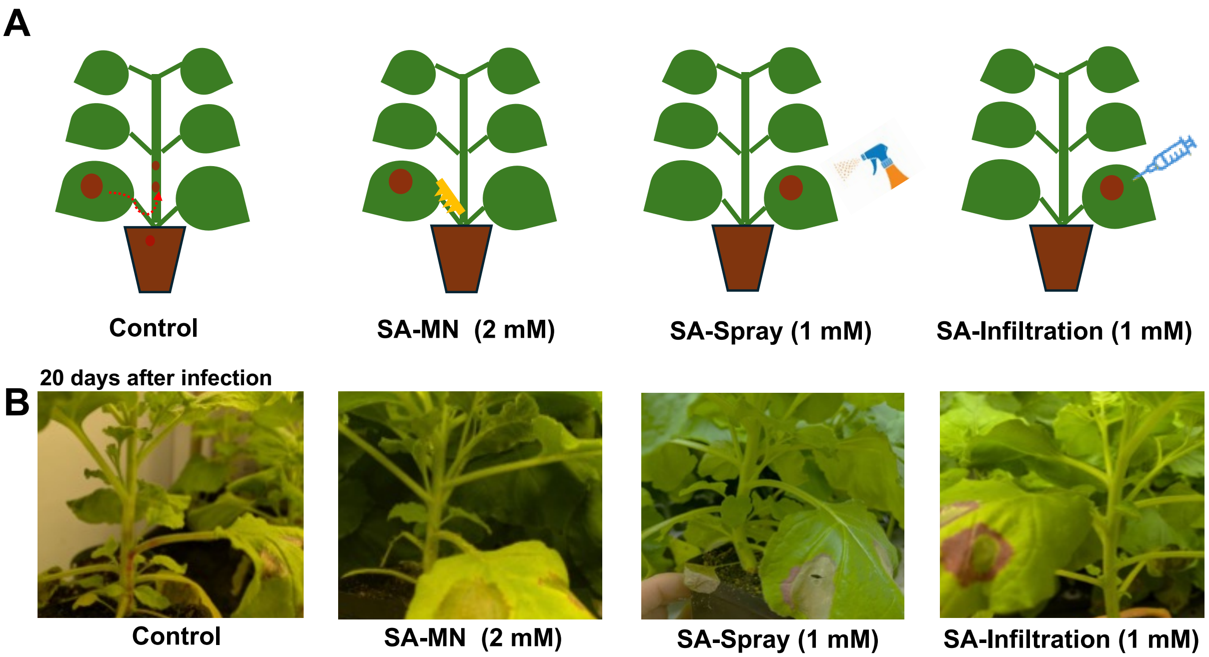


**Fig. S16: SA treatments inhibit TSWV movement from the leaf tissue to the stem in *Nicotiana benthamiana*, repeated with a higher SA dose in the spray and infiltration methods.** A) A model showing different SA delivery methods to *N. benthamiana*. B) Different phenotypes of TSWV infection. For the control plant, the TSWV movement as shown by red color can be observed on the stem after 20 days of infection, while all SA-treated plants remain healthy (no red color on the stem).

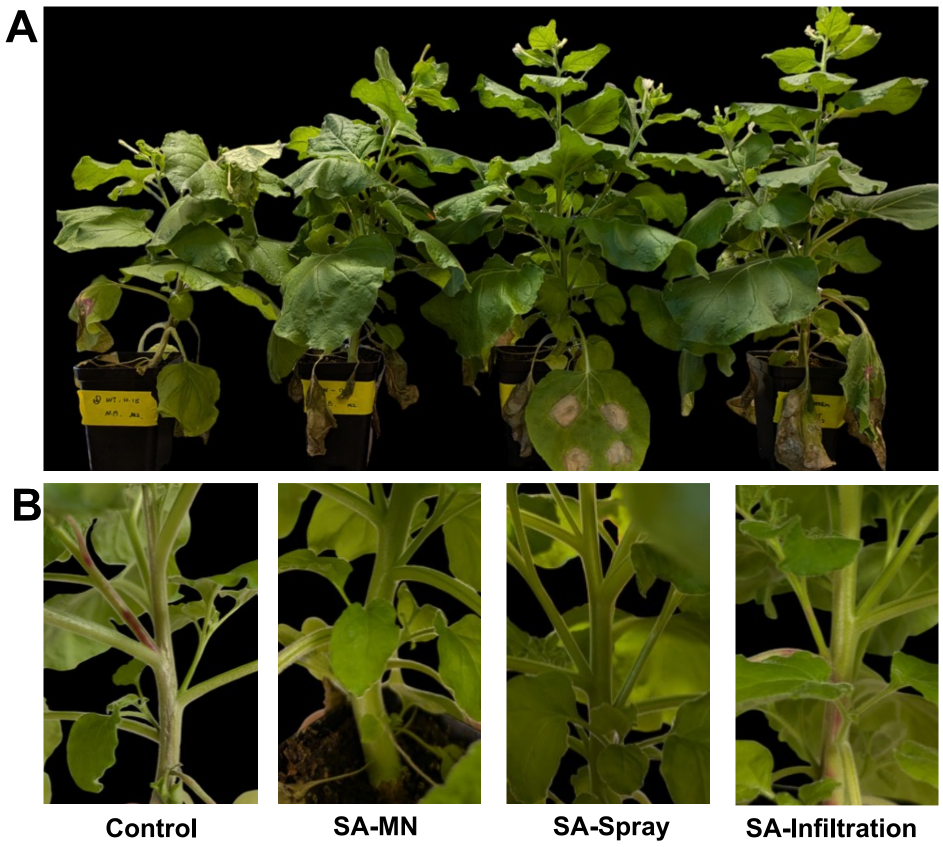


**Fig. S17: Phenotype of TSWV infection (30 days) in *Nicotiana benthamiana* with different SA treatments.** A) The phenotype of *N. Benthamiana* after TSWV infection, from left to right are Control, SA-MN, SA-spray and SA-Infiltration, respectively. B) Zoomed-in photos showing different phonotypes. In the Control plant, the stem necrotic symptoms appeared 30 days after infection. SA-MN and SA-Spray treated plants remained healthy, while SA-Infiltrated plants showed a red stem, indicating the presence of TSWV.

**Table S1. The primers used in the RT-qPCR analysis of the relative gene expression level in tomato and Arabidopsis**

| | |
|---|---|
| | tomato |
| *Slactin-F* | TGTGTTGGACTCTGGTGATGGTGT |
| *Slactin-R* | TCACGTCCCTGACAATTTCTCGCT |
| *SiGID1-F* | TCTTGTTGTTGTCGCAGGTT |
| *SiGID1-R* | CCCTATTGTTGCCTTCTCCA |
| *SiGID1b1-F* | GGCTGCTCTTCAATGGGTAA |
| *SiGID1b1-R* | TAAAACCTCGACGCCTGATT |
| | *Arabidopsis* |
| *AtUBC21-F* | CAGTGTTGCCAACGGTTCAT |
| *AtUBC21-R* | AAACGGAGGTATTCCGTAGTTGAG |
| *AtGID1a-F* | GATGTCTTGATTGATCGCAGGAT |
| *AtGID1a-R* | AGGAGGTTGCTCTTGATCTGCA |
| *AtGID1c-F* | AAGCAGGAAGAAGAACAGTGTAGTC |
| *AtGID1c-F* | AGTCAAAAGCTAACGCTAGAAAGC |
